# Chronic acoustic degradation via cochlear implants alters predictive processing of audiovisual speech

**DOI:** 10.64898/2026.01.25.701504

**Authors:** Simone Gastaldon, Flavia Gheller, Noemi Bonfiglio, Davide Brotto, Davide Bottari, Patrizia Trevisi, Alessandro Martini, Francesco Vespignani, Francesca Peressotti

## Abstract

This study provides the first neurophysiological evidence of how cochlear implant (CI) input affects predictive processing during audiovisual language comprehension in deaf individuals. Using EEG, we compared 18 CI users with 18 normal-hearing (NH) controls during sentence comprehension where final word predictability was determined by high or low semantic constraint (HC vs. LC) of the preceding sentence frame. Between sentence frame and final word, a 800 ms silent gap was introduced. Mouth visibility was manipulated during sentence frames (visible or digitally occluded; V+ vs. V-), while the final words were always presented with the mouth visible. In NH participants, lower-beta power (12-15 Hz) in left frontal and central sensors decreased for HC vs. LC contexts during the pre-target silent gap, but only when the mouths was visible, suggesting active prediction generation. In CI users, this lower beta power decrease was absent. After final word presentation, both groups showed N400 predictability effects, indicating preserved prediction evaluation. However, CI users exhibited extended N400 effects in the V+ condition, suggesting additional processing demands. Across all participants, pre-target beta modulations correlated with language production abilities, supporting prediction-by-production frameworks. Within CI users, poorer audiometric thresholds correlated with larger N400 constraint effects, possibly indicating greater reliance on contextual prediction to compensate for degraded sensory input. These findings demonstrate that CI-mediated perception alters the neural mechanisms of prediction generation. The link between production skills and predictive mechanisms suggests that strengthening expressive language abilities may enhance predictive processing in CI users.

## 1. Introduction

Language comprehension involves not only bottom-up decoding the incoming signal but also anticipating information at multiple representational levels. As language unfolds rapidly, listeners can form expectations about conceptual meaning, lexical semantics, syntactic structure, and even phonological features before such information becomes available in the input, with the aim of facilitating processing when that information is encountered (Heilbron et al., 2022; Huettig, 2015; Kuperberg & Jaeger, 2016; Kutas & Federmeier, 2011; Pickering & Gambi, 2018; Ryskin & Nieuwland, 2023). However, these mechanisms appear to be highly variable and flexible, and they seem to rely, at least in part, on additional cognitive resources, as suggested by findings from studies on aging, literacy level, and special populations (Favier et al., 2021; Federmeier et al., 2010; Gastaldon et al., 2024a; Jones et al., 2021; Milburn et al., 2023; Wlotko et al., 2012). Under this interpretation, predictive processing is not a single or uniformly deployed mechanism, but rather a set of strategies that individuals engage to different degrees depending on their sensory, cognitive, and experiential characteristics. For this reason, it becomes crucial to describe the “predictive profiles” of specific populations and to relate them to the cognitive and sensory processes that may be impaired, suboptimal, or differently organized (Gastaldon et al., 2024a). Identifying these profiles would allow researchers to gain a deeper understanding of predictive processing itself and, at the same time, to obtain a more complete picture of the linguistic and cognitive functioning of these populations. This line of work also has important applied value: it can inform scientific research, support clinical and rehabilitative practices, help communicate findings to educators and the general public, and ultimately contribute to more targeted and evidence-based approaches to assessment and intervention.

The present study focuses on one such population: people with profound deafness who use cochlear implants. Cochlear implants restore access to auditory information, but the signal they provide is spectrally degraded relative to natural hearing, with documented consequences for phonological processing, vocabulary development, and everyday communication (Hunter & Pisoni, 2021). Whether and how cochlear implant users engage in linguistic prediction, and whether compensatory mechanisms might support this process, remains poorly understood. This study aims to contribute to filling this gap.

### 1.1 Speech processing in cochlear implant users

Cochlear implants (CIs) are surgically implanted neuroprostheses that restore auditory perception by directly stimulating the auditory nerve. These devices are primarily implanted in cases of profound sensorineural deafness, in which the cochlea either is congenitally atypically developed or was damaged by traumatic injury or infection (Macherey & Carlyon, 2014). Among the genetic causes of sensorineural deafness, mutations in the GJB2 gene, which encodes Connexin-26 (Cx26), are a well-established marker. The Cx26 protein forms gap junctions between cochlear supporting cells, enabling intercellular transfer of ions and small metabolites that maintain the inner-ear metabolic homeostasis essential for normal hearing. Disrupted Cx26 production leads to inner-ear and cochlear dysfunction, resulting in impaired auditory encoding (Lefebvre et al., 2000). CIs have achieved remarkable clinical success in both congenitally deaf children and adults who lost hearing after language acquisition, with clinical assessments typically demonstrating impressive single-word and sentence recognition performance in quiet or moderately noisy conditions (Wilson & Dorman, 2008; Blamey et al., 2013). However, this clinical success masks a more complex reality: despite achieving high scores on standardized tests, many CI users report experiencing substantial effort and fatigue during everyday conversations (Hunter & Pisoni, 2021). Such disadvantages also appear to persist in individuals implanted very early in life (i.e., within the second year), suggesting that the development of normal-like spoken-language communication depends on a range of additional factors (Rinaldi et al., 2020). This discrepancy between laboratory performance and real-world communication challenges highlights a critical gap in our understanding of CI-mediated speech comprehension and underscores the profound impact that listening fatigue can have on social participation and quality of life (Philips et al., 2023).

The primary challenge confronting CI users stems from the fundamentally impoverished nature of the auditory signal, characterized by severely reduced spectral and temporal resolution (Hunter & Pisoni, 2021). While the healthy cochlea contains thousands of hair cells that provide frequency selectivity and preserve fine-grained temporal details, CIs bypass these damaged structures by delivering electrical pulses directly to the auditory nerve through an array of typically 12-22 electrodes (Wilson et al., 1991; Zeng et al., 2008). This dramatic reduction in independent information channels, effectively limited to 8-12 functional channels due to electrical current spread, means that the rich spectro-temporal details of natural hearing are compressed into a coarse representation. Modern devices employ sophisticated coding strategies that attempt to restore such temporal fine structure (Fischer et al., 2021). Yet, even with these advances, critical acoustic cues remain degraded: the spectral resolution necessary for distinguishing phonetically similar words is compromised, temporal fine structure that carries pitch and prosodic information is largely absent, and the dynamic range of natural hearing is compressed into a narrow electrical stimulation range (Zeng et al., 2008).

This degradation particularly impacts spoken-word recognition, a process that inherently involves temporal ambiguity as words unfold over time. During natural speech processing, early acoustic fragments are typically compatible with multiple lexical candidates. For instance, the onset /w/ could correspond to the beginning of the word “wizard,” “with,” “winner,” or “will.” In normal hearing (NH) listeners, graded activation is sustained over multiple phonological and lexical hypotheses that are rapidly pruned once disambiguating information is encountered, efficiently solving this transient ambiguity (Marslen-Wilson, 1987; Dahan & Gaskell, 2007; Gwilliams et al., 2018). With the spectrally degraded input conveyed by a CI, these same mechanisms are likely to operate over noisier and less informative evidence, making competition more prolonged (Friesen et al., 2001; Winn, 2016).

The cascade of processing challenges extends well beyond lexical recognition, reshaping the cognitive architecture that supports speech comprehension under degraded input. In NH listeners, limited fine-grained acoustic detail in the speech stimuli not only slows and destabilizes word recognition, but also makes it harder to build the rich contextual representations that normally scaffold predictive processing (Obleser & Kotz, 2010; Başkent et al., 2016). In CI users, this is compounded by increased listening effort: the same degraded input that motivates greater reliance on prediction also consumes the very resources needed to generate, maintain, and revise predictions, creating a potential vicious cycle in which effortful listening compromises higher-level language processing (Bhandari et al., 2021; Mattys et al., 2012). Converging physiological evidence supports this view. In a pupillometry study with postlingually deaf adult CI users and NH listeners hearing CI-simulated speech, Winn (2016) showed that semantic context produces a rapid “release” from listening effort in NH adults, but this anticipatory pupil reduction is delayed and attenuated in CI users and under vocoding, indicating slower or less efficient exploitation of predictive cues. Eye-tracking studies extend this picture to the time course of lexical competition. Adult CI users show broadly NH-like visual-world competition patterns, but with large individual differences: those with greater lexical uncertainty rely more heavily on sentential context to steer lexical access (Nagels et al., 2020). In younger populations, Children (5-10 year old) with CIs clearly use semantic prediction to facilitate spoken-word recognition, yet they are slower and less efficient than age-matched NH peers in suppressing phonological competitors (Blomquist et al., 2021). Further work with school-aged children with hearing loss (using hearing aids and/or CIs) shows that semantic prediction during sentence processing can be remarkably similar to NH controls (Holt et al., 2021), whereas syntactic prediction based on subject-verb agreement is present but systematically delayed in the same population (Davies et al., 2023). Altogether, these findings, largely from prelingually deaf, school-aged children with modern early devices on the one hand, and postlingually deaf adult CI users on the other, indicate that predictive processing is not globally absent under CI-mediated degradation. Rather, it tends to be slower, more resource-dependent, and more heterogeneous across individuals, with semantic versus syntactic prediction, and children versus adults, differentially affected depending on signal quality, linguistic representations, and the cognitive effort required to keep multiple hypotheses active over time.

Overall, rather than being uniformly absent or preserved, predictive mechanisms appear to be highly variable both within and across individuals, likely reflecting complex interactions between linguistic experience, available cognitive resources, and individual adaptation strategies to degraded auditory input. Moreover, substantial variability exists in device-related factors such as electrode placement precision, spiral ganglion cell survival, and the specific signal processing strategies implemented in each user’s device, introducing additional sources of individual differences that must be considered when interpreting variation in language processing abilities (Blamey et al., 2013). Understanding the sources and consequences of this variability is essential not only for advancing theoretical models of speech comprehension under adverse conditions but also for developing targeted interventions that leverage the flexible strategies CI users employ to optimize comprehension despite sensory limitations.

### 1.2 Multisensory speech: integrating visual mouth cues

Human speech perception, especially as part of face-to-face and situated communication, evolved as a fundamentally multimodal process (Benetti et al., 2023; Hagoort & Özyürek, 2025), with the visual system providing complementary information that becomes increasingly valuable when auditory input is compromised. The integration of auditory and visual speech information represents a sophisticated computational achievement, allowing listeners to extract linguistic content from correlated patterns across sensory modalities, most notably the temporal synchrony between acoustic events and visible body cues of the interlocutor. Such visual cues include not only articulatory mouth movements, but also the hands, head, and upper body. Lip and jaw movements provide fine-grained segmental and prosodic information, while co-speech hand gestures, either conveying iconic meaning or providing a rhythmic anchoring, supply additional (often predictive) constraints on what is likely to be said next. Corpus and experimental work show that meaningful hand gestures typically start before the corresponding spoken information, giving listeners advanced access to semantic content and enabling multimodal prediction (Holler & Levinson, 2019; ter Bekke et al., 2024). In adverse or CI-like listening conditions, such as vocoded speech that mimics the limited spectral resolution of implants, beat and iconic gestures jointly with visible speech enhance comprehension and can even bias lexical stress perception, acting as a kind of “manual McGurk” cue that helps listeners disambiguate prosodic structure (Bosker & Peeters, 2021; Drijvers & Özyürek, 2017; Maran et al., 2025). Hearing-impaired listeners and CI users appear particularly well placed to exploit these multimodal affordances (Obermeier et al., 2012; Sparrow et al., 2020).

In visual articulatory cues in particular, the dynamic movements of the lips, jaw, and tongue provide fine-grained information about place and manner of articulation, and even aspects of prosodic structure, thereby constraining the set of viable phonetic and lexical candidates and reducing the uncertainty inherent in degraded or noisy auditory signals (Peelle & Sommers, 2015; Stacey et al., 2016). This audiovisual benefit extends well beyond simple redundancy. Visible speech can disambiguate acoustically confusable phonemes, provide advance information about upcoming articulatory events, and sharpen the temporal segmentation of continuous speech, all of which become critical when the auditory channel is compromised (Benard & Başkent, 2015; Blackburn et al., 2019; Sumby & Pollack, 1954). In CI users, evidence for enhanced audiovisual integration is both robust and multifaceted, revealing fundamental reorganization of multisensory processing following auditory deprivation and subsequent restoration via the implant. Behavioral and neuroimaging work consistently shows that CI users rely more on visual speech cues and derive larger relative benefits from audiovisual input than NH listeners, a pattern reported across postlingually deaf adults and, more recently, in children with CIs as well (Strelnikov et al., 2009, 2015; Stevenson et al., 2017; Alemi et al., 2023). This enhanced visual reliance transcends simple improvements in lipreading ability and is accompanied by characteristic changes in brain organization: occipital and posterior temporal regions involved in visual and audiovisual processing show increased engagement, and their activity predicts later auditory recovery after implantation, indicating crossmodal plasticity in service of speech comprehension (Strelnikov et al., 2013, 2015). In addition, CI users frequently exhibit superadditive integration effects, where audiovisual performance exceeds what would be predicted from the sum of auditory-only and visual-only performance, pointing to qualitative changes in multisensory integration mechanisms rather than mere quantitative shifts in sensory weighting (Stevenson et al., 2017; Moberly et al., 2020). In McGurk-type paradigms and audiovisual speech identification, CI users often show stronger use of visual speech than NH controls, suggesting that long-term experience with degraded auditory input drives compensatory enhancements in crossmodal processing (Rouger et al., 2008; Schorr et al., 2005; Strelnikov et al., 2015).

Crucially, the magnitude of audiovisual benefit in CI users is strongly modulated by linguistic structure and predictability: benefits tend to be modest for isolated phonemes or nonsense syllables but increase markedly for words and are maximal for sentences in which contextual constraints can guide interpretation (Strelnikov et al., 2009; Stacey et al., 2016; Choi et al., 2024). This pattern suggests that visual speech does not merely supplement a noisy auditory stream through bottom-up sensory fusion. Instead, it actively interacts with top-down predictive mechanisms that exploit linguistic context to pre-activate likely upcoming segments and words (Peelle & Sommers, 2015). Both prediction and audiovisual integration serve to reduce uncertainty, prediction by constraining the hypothesis space based on context, and visual speech by providing complementary, time-locked evidence about articulatory events. Their convergence implies that CI users’ enhanced reliance on visual articulatory cues may fundamentally alter not only what information is available for comprehension, but also how predictive language processing is implemented under conditions of chronic sensory limitation.

### 1.3 The present study

Despite extensive research documenting both predictive processing differences and enhanced audiovisual integration in CI users, a critical gap remains in our understanding of how these two fundamental aspects of language comprehension interact. The existing literature has largely examined prediction and multisensory integration as independent phenomena, with studies of predictive processing typically employing auditory-only or written stimuli that preclude investigation of audiovisual interactions, while research on audiovisual integration has focused primarily on perceptual outcomes without examining underlying predictive mechanisms. This methodological separation has left unresolved the crucial question of whether and how visual speech information modulates the anticipatory processes that support efficient language comprehension. Recent evidence hints at potential interactions: neural oscillatory responses show stronger modulation by sentence-level constraints during audiovisual compared to auditory-only speech processing (Brunellière et al., 2022), though these effects have been interpreted primarily through the lens of enhanced attentional engagement rather than as evidence for visual modulation of predictive processing per se. The absence of studies directly examining how visual speech cues shape the generation and evaluation of linguistic predictions represents a significant theoretical gap, particularly given that both processes fundamentally serve to reduce uncertainty during comprehension.

Most critically, to the best of our knowledge, no study to date has employed electroencephalography (EEG) to investigate the neural correlates of predictive processing in CI users, leaving us without crucial neurophysiological evidence about how degraded auditory input and compensatory visual reliance jointly influence the temporal dynamics of language comprehension. This gap is particularly consequential given that EEG offers insights into the cascade of predictive and integrative processes that unfold during real-time language processing, processes that behavioral measures alone cannot fully capture. The absence of such neural evidence prevents us from understanding whether CI users’ documented behavioral differences in prediction reflect fundamental alterations in predictive mechanisms, strategic adaptations in how predictions are deployed, or differences in the integration of predicted and observed input. Furthermore, without examining these processes in naturalistic audiovisual contexts that reflect how CI users actually experience speech, we risk overlooking adaptive strategies that may only emerge when multiple information sources are available. Given CI users’ demonstrated enhancement in audiovisual integration and the theoretical importance of understanding how sensory and cognitive adaptations interact, investigating the neural basis of prediction under audiovisual conditions is essential for developing comprehensive models of language processing following sensory restoration and for optimizing clinical interventions that leverage these adaptive mechanisms. Here, we use a widely adopted paradigm in which participants hear sentence frames (sentences missing the final word) followed by a brief silent gap before the target. This design is well suited to probing anticipatory/predictive processing, since the semantic constraint of the sentence frame can either strongly narrow down the upcoming target (high constraint) or leave multiple plausible endings (low constraint). By doing so, it selectively modulates the extent to which participants can generate specific predictions, and therefore the neural dynamics expressed during the silent pre-target gap. Moreover, depending on the type of target (e.g., a picture to name or judge, or an auditory word to integrate), the same framework can be used to investigate predictive mechanisms in both language production and comprehension (Gastaldon et al., 2020, 2023, 2024b; Hustá et al., 2021; León-Cabrera et al., 2022; Piai et al., 2014, 2015, 2016, 2018; Roos & Piai, 2020; Roos et al., 2024). In the present study, we implemented a comprehension-focused version of the paradigm using audiovisual materials: sentence frames and target words were presented as videoclips of a speaker, and participants were asked to listen to the sentences.

## 2. Materials and Methods

All procedures performed in this study adhered to the principles of the Declaration of Helsinki. Ethical approval was granted by the Ethical Committee for Psychological Research of the University of Padova (protocol n. 4188). Participants received detailed information about the study and provided written informed consent. In the case of minors, written informed consent was obtained from both legal guardians and minor. Data was collected, stored, and processed in compliance with current regulations on privacy and research ethics.

### 2.1 Participants

CI users (N = 18; 10 females, 7 males, 1 preferred not to declare) were recruited on a voluntary basis through the Otorhinolaryngology Unit of Padova University Hospital. To be eligible, CI users had to be at least 12 years old, have a diagnosis of profound deafness (either congenital, progressive or due to traumatic injury), and have undergone cochlear implantation. Participants were required to have no history of psychiatric disorders, developmental comorbidities, or other diagnosed sensory, motor, or cognitive impairments. Given the challenges in recruiting this population with such restrictive criteria, sample size was not predetermined, either through power analysis or other methods. Instead, the final sample size reflects the most feasible compromise between time constraints and the limited availability of participants meeting the inclusion criteria and willing to participate (Lakens, 2022). For privacy and data-protection reasons, demographic and clinical information are not reported in a participant-level table, as combinations of variables in a small sample could enable re-identification. Instead, we describe the sample in aggregated form (Figure 1), consistent with GDPR principles of data minimization and privacy (Regulation (EU) 2016/679, Art. 5 and Art. 25) and the focus on identifiability (Recital 26).

**Figure 1.**
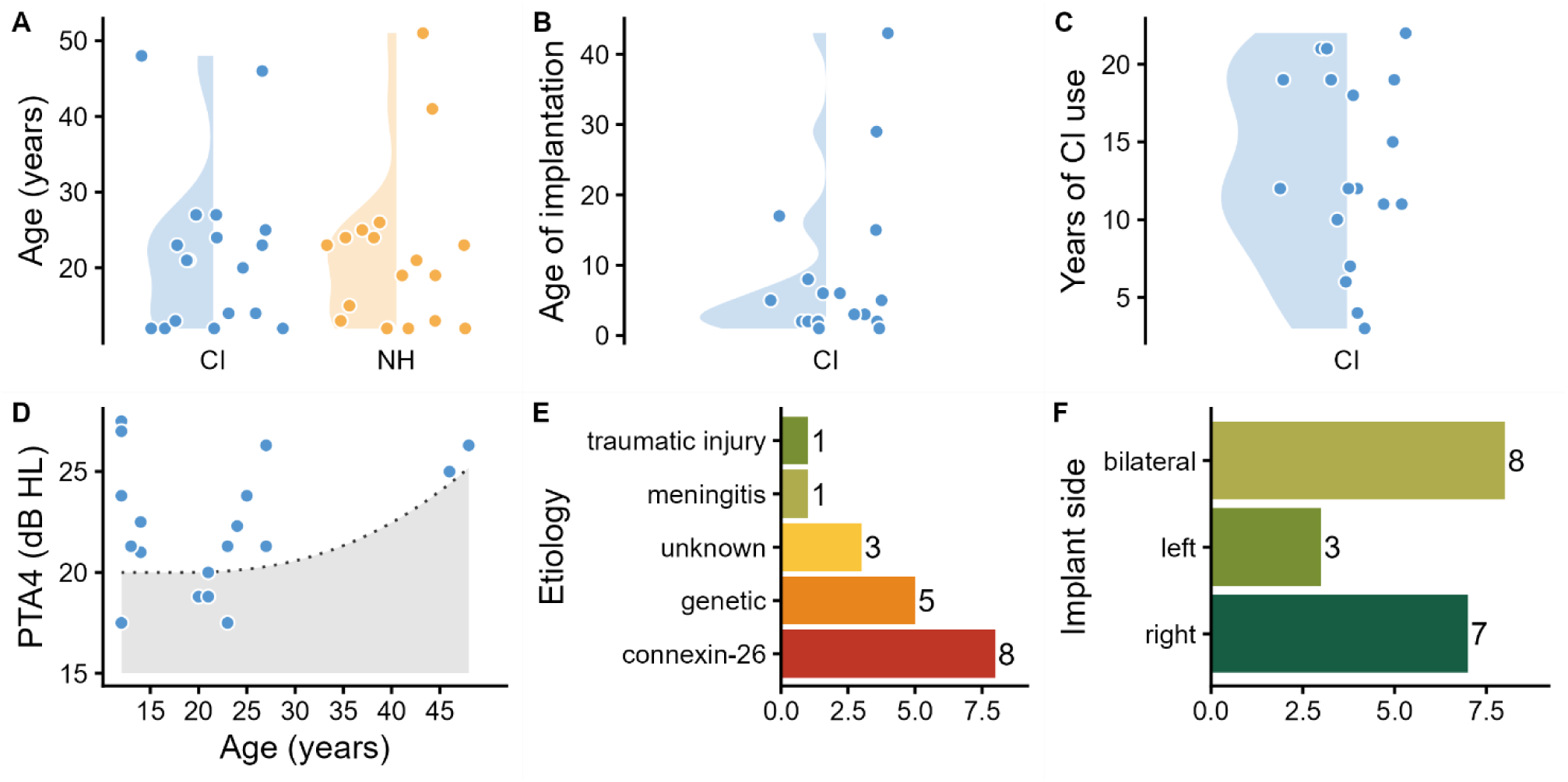
Experimental sample. Panel A) Distributions of age across CI users and NH controls. Panel B) Age of implantation in CI users. Panel C) Number of years of CI use. Panel D) Individual PTA4 thresholds plotted against age with an age-adjusted normative threshold from ISO 7029:2017. The grey area represents normal hearing (upper limit = median + ∼20 dB). For ages below 18, the threshold is fixed at 20 dB HL, as ISO 7029:2017 covers only ages 18-80. Points above the grey shaded area indicate thresholds that exceed age-appropriate norms for normal-hearing individuals. The figure is intended to be purely descriptive, illustrating that overall thresholds are generally higher than normative values, with no evident association with age. This information was retrieved from the participant’s most recent audiological assessment. Panel E) Etiology for sensorineural deafness. When the etiology is reported as ‘genetic’, this indicates a supposedly congenital origin in the absence of positive tests for mutations in the GJB2 gene encoding the protein Connexin-26. Panel F) Side of implantation.

Across the 18 CI users who took part in the study, implantation patterns were mostly bilateral (N = 8), primarily sequential rather than simultaneous, and unilateral with right-ear dominance (N = 7). Ages ranged widely (12-48 years), but most were adolescents (12–14-year-old; N = 7) or young adults (20–27-year-old; N = 9), with two older adults (48– and 46-year-old). Age of implantation was typically very early, with more than half of the sample implanted between 1-5 years of age. Accordingly, years of device use spanned 3-22 years. From an audiological perspective, participants showed variable implant-aided audibility. PTA4 (Pure Tone Average 4; mean threshold in dB across 500, 1000, 2000, and 4000 Hz) ranged from 17.5 to 27.5 dB, indicating substantial inter-individual variability. Most participants exceeded 20 dB, a commonly used upper limit for typical hearing in young adults. The predominant etiology was Connexin-26-related deafness (N = 8), followed by other non-further specified genetic causes (N = 5), with isolated cases of meningitis (N = 1), traumatic injury (N = 1), and unknown specific origin (N = 3). All participants acquired spoken Italian as their first language, including prelingual deaf participants implanted early in life, who were raised in Italian speaking households. None of the participants and their families reported knowledge and use of Italian Sign Language (LIS). In fact, in Italy there is a tendency for implanted individuals not to learn sign language alongside spoken language, unless they are children of deaf parents (Rinaldi et al, 2020).

A group of NH participants (N = 18; 11 females, 7 males) was also recruited using a snowball procedure. Manual preference was assessed for both groups with an Italian adaptation of the Edinburgh Handedness Inventory, long version (Oldfield, 1971). The groups did not significantly differ for age (CI: mean = 21.89 years, SD = 10.66; NH: mean = 21.39, SD = 10.46; t = −1.19, p = 0.89) and handedness laterality index (CI: mean = 80.5, SD = 38.66; NH: mean = 83.52, SD = 27.26; t = −0.27; p = 0.79). Participants were given a compensation of 25€ for their participation for the overall testing sessions (language skills and EEG session; see below).

### 2.2 Language skills assessment

In addition to the main experiment, we administered a set of tasks to assess individual linguistic skills. These tasks provided both a general evaluation of language abilities in CI users and NH controls, and data for correlational analyses with electrophysiological markers of predictive processing. Two tasks targeted production abilities, while two assessed comprehension abilities.

For the production assessment, participants completed verbal fluency tasks (semantic and phonological) and a sentence generation task, administered in accordance with the guidelines of the BVN 12-28 (Gugliotta et al., 2009), a standardized neuropsychological battery designed for individuals aged 12 to 18 years. This choice was motivated by the expectation of recruiting a large proportion of underaged participants, as the annual number of cochlear implant procedures in Italy has more than doubled over the last 20 years, according to the Italian Registry of Auditory Implantable Devices (RIDIU)^1^. In fact, of the 18 CI users we recruited, 7 are under 18 years of age. While the battery norms are based on adolescents, the test of sentence generation remains useful for describing linguistic performance, with no clinical or neuropsychological assessment intent. Verbal fluency was assessed using both semantic and phonemic fluency tasks. In each task, participants were given one minute to produce as many words as possible according to the specified criterion. For semantic fluency, participants were asked to generate words belonging to the categories animals and fruit. For phonemic fluency, they were asked to generate words beginning with the phonemes /f/ and /l/. Responses were recorded using Audacity on a laptop positioned in front of the participant. Recordings were manually started and stopped by an experimenter for each trial. This procedure yielded four recordings per participant, corresponding to the four fluency trials (two semantic and two phonemic). For each cue (“animal”, “fruit”, /f/, /l/), the number of correct words produced was calculated. From the list of produced words, for each participant, were excluded: 1) repeated words, 2) variations of the word said immediately before (e.g., *cervo/cerbiatto*, deer/fawn*, figlio/figlia*, son/daughter), 3) words composed from immediately preceding words (e.g., *pesce/pesce palla*, fish/pufferfish or blowfish]). Then, for each participant we averaged the number of correct words relative to the semantic and phonological cues, obtaining two fluency measures, semantic and phonological, respectively. In the sentence generation task, participants were instructed to produce a meaningful sentence that included a pair of nouns read aloud by the experimenter. They were asked not to use the conjunction e (“and” in Italian) and were given unlimited time to respond. Five noun pairs were selected from the BVN test battery, and an additional pair (*matita-carta*, pencil-paper) was presented as an example before testing. The selected pairs were *filo-bottone* (thread-button), *fuoco-legna* (fire-wood), *casa-luce* (house-light), *sciopero-salario* (strike-salary), and *vita-libertà* (lifefreedom). All responses for this task were recorded in a single Audacity file. Offline, a score was attributed to each sentence according to the guidelines of the BVN 12-18 (Gugliotta et al., 2009), then the sum was taken as a global score for each participant. Each sentence was given a score ranging from 0 to 3 according to semantic appropriateness and syntactic and grammatical correctness.

For the comprehension assessment, participants completed computerized tasks targeting complementary aspects of lexical and grammatical processing. The lexical decision (LD) task evaluates the efficiency of lexical access, requiring participants to rapidly discriminate real words from phonotactically plausible non-words; this taps into stored lexical representations and word-form processing. In contrast, the sentence-picture matching (SPM) task assesses sentence-level comprehension, requiring participants to integrate lexical information with syntactic structure and thematic roles to select the picture that matches the meaning of the sentence. Together, these tasks provide a broad characterization of receptive linguistic abilities, covering both single-word processing and the ability to compute sentence meaning. Participants were asked to read the instructions and to ask for clarification if needed. For LD, we used the stimuli in Amenta et al. (2021), consisting of 90 items (45 words and 45 non-words). The task was delivered through PsychoPy2 (Peirce et al., 2019). Participants were presented with words and phonotactically legal non-words in written form and were required to press two different keys on the keyboard to categorize the string as an Italian word or not. Each string was preceded by a fixation cross (500 ms) and remained on the screen until response. Reaction times (RTs) and accuracy were recorded. For SPM, participants were shown a written sentence at the centre of the screen for 2 seconds. Subsequently, four pictures appeared on four sections of the screen (upper left, upper right, bottom left and bottom right), which remained until a response was provided. For each sentence, participants had to use the mouse to select the correct illustration from four options: a target picture, two distractors mismatching either the characters or the action, and a fully unrelated distractor. RTs and accuracy were recorded. The task was administered in OpenSesame (Mathôt et al., 2012), using the materials and implementation provided by Artesini (2019). In Artesini (2019), the images were adapted from the *Comprendo* test (Cecchetto et al., 2012) and paired with sentences specifically created for Artesini’s work. For the present study, to keep the session within a manageable duration, we selected a subset of 40 sentences distributed across three categories: 8 active sentences, 16 passive sentences, and 16 sentences with clitic pronouns (see Supplementary Material on OSF for further details). Importantly, this task evaluates not only knowledge of grammatical rules but the ability to integrate them into meaningful sentence interpretation, skills that may dissociate, as individuals can show intact grammar knowledge while still failing to understand sentence meaning.

### 2.3 Audiovisual speech comprehension: stimuli

A subset of sentences was selected from the stimuli used in Gastaldon et al. (2020, 2023, 2024b). First, an online cloze test was administered using the original, validated full set of stimuli, specifically targeting participants under 18 years of age. This choice was motivated by the need to assess whether all highly constraining sentences remained valid for a younger population, as some might have lost their predictive value due to cultural changes over time. Moreover, this procedure reduced the total number of stimuli and consequently the testing time, which was particularly important given that pre-adolescents, especially those in the CI group, were expected to experience fatigue and diminishing attention earlier than young adults. Ten respondents participated in the questionnaire (mean age = 14.6 years, SD = 1.9). For the final set of sentences, we selected only those with a high cloze probability (CP) in the high-constraining (HC) condition and no words with a moderate cloze probability in the low-constraining (LC) condition. This selection resulted in a total of 196 sentences, 98 HC (mean CP = 0.9, SD = 0.11) and 98 LC (mean CP = 0.04, SD = 0.07), with statistically significant difference between constraints, assessed by means of Welch’s independent sample t-test (t = 66.34, df = 169.37, p < 0.001). Each target word was thus associated with one HC and one LC sentence. Subsequently, a young female native speaker of Italian was video recorded while uttering the sentences and target words separately. The recording setup included a video camera positioned three meters from the speaker and two microphones placed on a table in front of the speaker. The microphones and video camera were connected to a mixer, producing an integrated audio–video recording (MP4 file) for each sentence and each target word. The videos were recorded at 30 frames per second, with audio sampled at 48 kHz. The audio-video files were then cropped using the software Kdenlive so that the speaker’s nose was centered in the frame, resulting in a final resolution of 800 × 720 pixels. For each sentence file, an additional version was created in which the mouth was obscured by a grey rectangle covering the area from just above the upper lip to the chin. Target files were not manipulated in this respect, so that during the experiment, mouth visibility is manipulated only during the sentence frame. Stimuli were then divided into two sets, A (HC: mean = 0.91, SD = 0.107; LC: mean = 0.045, SD = 0.068) and B (HC: mean = 0.894, SD = 0.113; LC: mean = 0.037, SD = 0.076). In these lists, each pair of sentence frames and their target word were either assigned to the V+ or the V-condition (in this way the target word appears twice within the same mouth visibility condition), so that across the experiment all stimuli were presented in the mouth covered and mouth visible condition. Stimuli are available in the OSF repository (see Data availability statement).

### 2.4 Audiovisual speech comprehension: procedure

Participants performed the comprehension task in a dimly lit, sound-attenuated room while EEG was continuously recorded. Stimuli were presented on a computer screen positioned approximately 70 cm from the participant. Audio was delivered through two loudspeakers placed symmetrically on each side of the screen. Because of expected individual variability in auditory thresholds, sound intensity was adjusted to a comfortable level for each participant.

Prior to the main experiment, participants completed a short practice session to familiarize themselves with the task; none of the practice items appeared in the experimental set. In each trial, participants watched a video of a native Italian speaker producing a sentence. Trials began with a blank placeholder (1 s), followed by the sentence frame (the sentence up to, but not including, the final word). Before the presentation of the target word, a silent 800 ms gap was introduced, during which a blank placeholder was displayed. Mouth visibility during the target frame was not manipulated and was always visible. The decision to manipulate mouth visibility only during sentence frames was guided by the rationale that predictions are generated during and after processing the frame, and the aim of the present study was to observe whether masking articulatory cues could affect this ability, all being equal at the target presentation. To ensure attention throughout the task, comprehension yes/no questions were presented at the end of 20% of the trials. The entire task lasted approximately 40 minutes, with short breaks every 28 trials. A pseudorandomized list was generated for each participant by using Mix (van Casteren & Davis, 2006), so that each target word was presented the second time at least 7 trials after the first presentation, the same constraint and mouth visibility condition was presented for maximum 3 consecutive trials. The task structure is shown in Figure 2.

**Figure 2.**
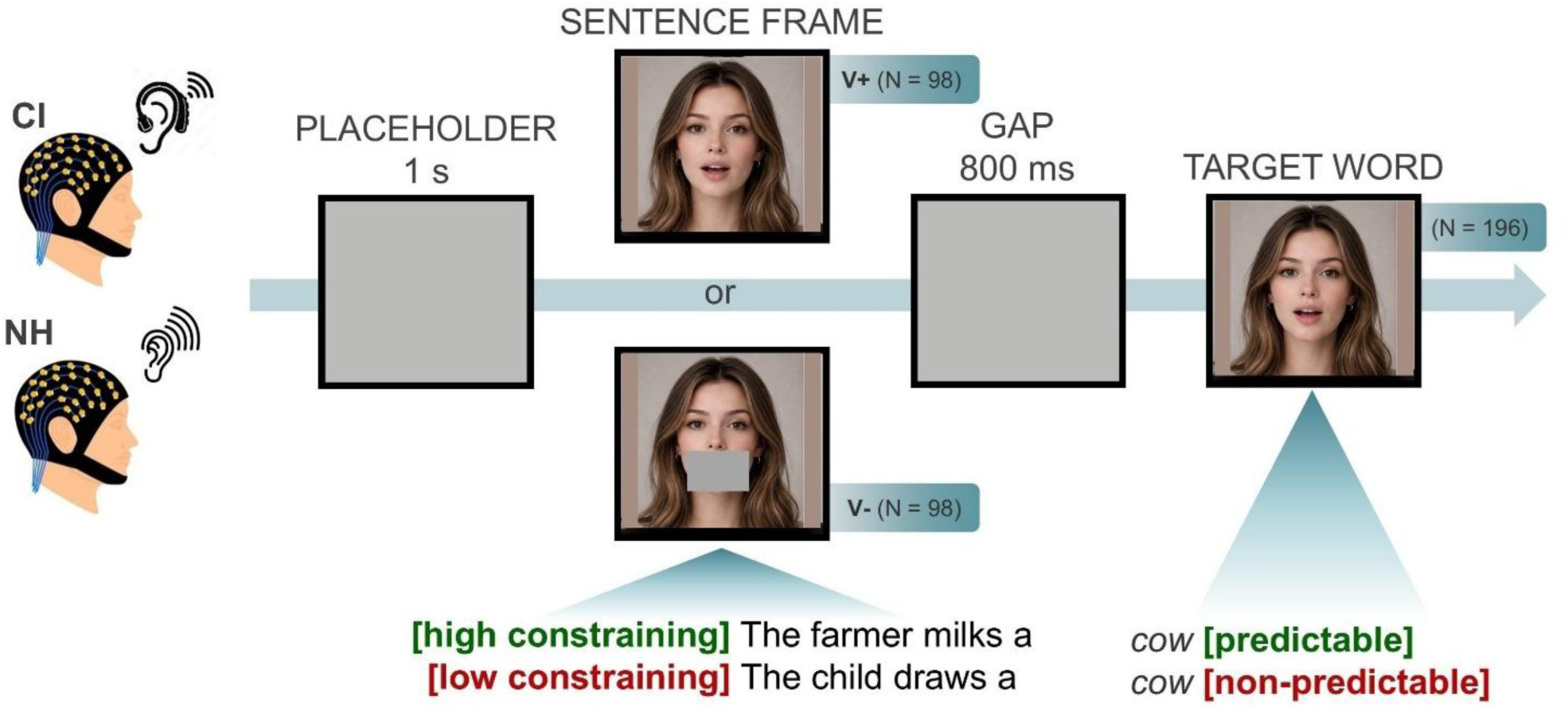
Trial structure of the audiovisual sentence comprehension task. According to the set (A/B) assigned to the participant, the pairs of sentence frames associated with the target could either be in the V+ or V-condition. Mouth visibility conditions (V+/−) were interspersed throughout the experiment and not presented in separate blocks.

### 2.5 EEG recording and preprocessing

EEG data were recorded using a BrainProducts BrainAmp system with an EasyCap montage with 64 active Ag/AgCl electrodes positioned according to the 10-20 system. Electrode locations were assigned using the Colin27 BrainProducts EasyCap M1 template to ensure standardized spatial coordinates across all participants. Sixtyone electrodes were used as EEG channels, covering frontal left (n = 16; Fp1, Fp2, AF7, AF3, AFz, AF4, AF8, F7, F5, F3, F1, Fz, F2, F4, F6, F8), frontal right (n = 9; FT9, FT7, FC5, FC3, FC1, FC2, FC4, FC6, FT10), central (n = 9; T7, C5, C3, C1, Cz, C2, C4, C6, T8), centro-parietal (n = 9; TP7, CP5, CP3, CP1, CPz, CP2, CP4, CP6, TP8), parietal (n = 9; P7, P5, P3, P1, Pz, P2, P4, P6, P8), parieto-occipital (n = 5; PO7, PO3, POz, PO4, PO8) and occipital (n = 3; O1, Oz, O2) scalp areas. Three electrodes were used to record electro-oculographic activity: 2 for horizontal eye movements, placed at the outer canthi (TP9 as HEOG_left and TP10 as HEOG_right) and one for vertical movements placed below the left eye (Iz as VEOG_down). For ground and reference electrodes, the standard positions of the cap were used (ground between Fp1 and Fp2; reference at FCz).

Data preprocessing was performed with Brainstorm (Tadel et al., 2011). Raw data were first high-pass filtered to remove slow drift and DC components, with 0.5 Hz as cut-off frequency and an attenuation of 60 dB. One CI participant was filtered at 1 Hz due to slow drifts surviving the 0.5 Hz cut-off filtering. However, such slow drifts were still present throughout the recording after filtering, thus this participant was not included in the ERP analyses to avoid distortions of ERP components. However, this participant was maintained for time-frequency analysis as filtering at 1 Hz does not impact decomposition at the frequencies of interest (7-30 Hz) and drifts were not affecting the range of interest. Bad channels and bad recording segments were marked manually. Note that, in the CI group, recording from temporo-parietal electrodes over the hemisphere corresponding to the implanted side was not possible because of external implant components. To avoid damaging the device, no conductive gel was applied at these sites, and the corresponding channels were marked as bad for subsequent analyses. As a result, the CI group had a higher number of bad channels than the NH group (Supplementary Fig. S1). Therefore, group comparisons involving temporo-parietal sensors were not meaningful and were not performed.

Independent Component Analysis (ICA) decomposition was employed to remove ocular artifacts (blink, saccades) from continuous EEG data. ICA (Infomax algorithm) was applied to a band-passed filtered copy of the data (2-100 Hz) resampled to 500 Hz to optimize computational efficiency and artifact identification. The number of ICA components was calculated individually for each participant using the formula: 61 – (number of bad channels + 5), accounting for the total electrode montage and individual channel quality. Components were automatically sorted based on their correlation with EOG signals to facilitate identification of artifact-related components. ICA components were manually examined and excluded when topography and time-series described clear EOG artifacts and possible non-physiological channel-specific artifacts. The average number of rejected IC was 4.5 (SD = 1.98) for the CI group, and 3.22 (SD = 1.17) for the NH group.

After ICA cleaning, additional artifact detection was performed to identify residual slow-wave artifacts in the 1-7 Hz frequency range using a conservative threshold of 5 SDs. Detected segments were marked as ‘bad’ for subsequent exclusion during epoching. Finally, because of the centroparietal distribution of the N400, one of the electrophysiological markers of interest, the EEG data was re-referenced more anteriorly to the AFz electrode and therefore enhance the N400 detectability. This choice also ensures that reference is as far as possible from the implant with no lateralization biases.

#### 2.5.1 Time-Frequency decomposition

Time-frequency decomposition was performed using Morlet wavelets to examine oscillatory activity in response to gap stimuli normalized against activity prior to sentence onset. Prior to wavelet analysis, data were epoched with extended time windows (–1 to 4 seconds for experimental conditions, time-locked at gap onset, and –1 to 1.5 seconds for baseline, time-locked at sentence placeholder onset). Epochs were bandpass filtered between 2-80 Hz to facilitate artifact detection. Automatic trial rejection was implemented using a peak-to-peak amplitude criterion (200 μV peak-to-peak in any EEG channel), with condition-specific time windows for artifact detection (–0.5 to 0.5 seconds for baseline, –0.3 to 1.5 seconds for gap conditions). In the CI group, the mean number of epochs retained for time–frequency analysis was 184.3 (SD = 8.96; 94.1%) for the baseline condition, and 43.8, 44.4, 43.8, and 44.7 for the HC V+, LC V+, HC V-, and LC V-conditions, respectively. These corresponded to retention rates of 89.3%, 90.6%, 89.3%, and 91.2%, with standard deviations between 3.6 and 5.2 epochs, indicating stable data quality across conditions. In the NH group, baseline retention was similarly high, with a mean of 185.5 epochs (SD = 20.7; 94.6%). For task-related conditions, the number of retained epochs was 45.4 for HC V+, 45.6 for LC V+, 44.8 for HC V-, and 46.1 for LC V-, corresponding to retention rates of 92.7%, 93.0%, 91.5%, and 94.0%, respectively (SDs = 4.0–5.3). Overall, both groups showed high and comparable retention rates across baseline and experimental conditions, confirming the robustness and balance of the dataset for subsequent time–frequency analyses. For each retained epoch, Morlet wavelet decomposition was computed for frequencies between 8-30 Hz using a central frequency of 1 and a full-width at half-maximum temporal resolution (FWHM) of 3. Baseline activity was calculated as the average in the interval from 0 to 0.3 seconds, averaging across all experimental conditions of the baseline epochs (condition-average baseline; Cohen, 2014). Event-Related Spectral Perturbation (ERSP) values separately for each condition were calculated by normalizing gap condition power relative to baseline power using the formula: ERSP = (Powergap – Powerbaseline) / Powerbaseline × 100. The resulting modulation is therefore to be interpreted as % change relative to the processing of the placeholder at the beginning of the trial (empty rectangle before the video of the sentence frame, hence a visual stimulus in absence of auditory/speech information). Final time-frequency maps were extracted for the analysis window of –0.5 to 3 seconds relative to gap onset. Finally, for computational limitations, condition-averaged time-frequency matrices for each participant were resampled to 250 Hz (i.e., a temporal resolution of 4 ms) and subsequently exported for statistical analysis.

#### 2.5.2 Event-Related Potentials

For all ERP analyses, the participant CI_01 was excluded to avoid distorting components of interest due to intensive slow drifts which were still visible after high-pass filtering at 1 Hz, a threshold that is considered to introduce distortions (Tanner et al., 2015). Continuous data were epoched around target onsets with a time window of –0.2 to 2 seconds. Baseline correction was applied using the pre-stimulus interval (–0.2 to 0 seconds). Epoched data were low-pass filtered at 20 Hz using a strict 60 dB attenuation filter to remove high-frequency noise while preserving ERP components of interest (Zhang et al, 2024). Automatic trial rejection was implemented using a peak-to-peak amplitude criterion, with trials exceeding 150 μV in any EEG channel being automatically excluded from further analysis. This threshold was chosen to remove trials contaminated by residual artifacts while preserving sufficient trial counts for reliable averaging. In the CI group, the mean number of retained epochs was comparable across conditions: 40.6 for HC V+, 41.0 for LC V+, 40.8 for HC V-, and 41.8 for LC V-. These values correspond to retention rates of 82.8%, 83.7%, 83.3%, and 85.2%, respectively. Variability across participants ranged from about 6 to 8 trials (SD = 5.9–8.0). In the NH group, retention was slightly higher and more stable across conditions: 44.7 for HC V+, 43.6 for LC V+, 44.1 for HC V-, and 44.8 for LC V-, corresponding to 91.2%, 88.9%, 89.9%, and 91.5% of total trials retained. Standard deviations ranged from 3.7 to 5.3 trials. Overall, epoch retention was high and balanced across all conditions, ensuring comparable data quality between groups and conditions for subsequent analyses. Individual condition averages were computed for each participant using arithmetic mean across accepted trials. ERPs were calculated separately for each of the four experimental conditions for each participant and subsequently exported for statistical analysis (sampling rate of 1000 Hz, i.e., a temporal resolution of 1 ms).

### 2.6 Statistical analyses

All statistical analyses were conducted with R (v 4.3.3) within the RStudio environment (v 2023.06.1). In the following, we describe the rationale for the analytical approach adopted for each of the measures presented in the study.

#### 2.6.1 Language skills

Response times (RTs) and accuracy of both SPM and lexical decision tasks were analyzed by means of generalized linear mixed-effects models (all mixed models in the study were implemented with the R package *lme4*, Bates et al., 2015; to estimate p-values we used the package *lmerTest*, Kuznetsova et al., 2017). Specifically, RTs were modeled with a Gamma distribution family and log link function for skewed data with positive values, while accuracy was modeled with a binomial distribution. All categorical predictors were coded with sum contrasts so that main effects reflect deviations from the overall mean. In the production tasks, fluency (total number of words produced) and sentence generation scores (scored by following the guidelines in Gugliotta et al., 2009) were tested by means of Mann-Whitney U test (Wilcoxon rank-sum test). Effect sizes are calculated as r (rank-biserial correlation), a measure of the strength and direction of the difference between the two groups.

#### 2.6.2 Time-frequency representations

Six regions of interest (ROIs) were defined based on selected electrodes to cover functionally relevant scalp areas (Supplementary Figure S1). The central ROI included C1, Cz, C2, CP1, CPz, CP2, P1, Pz, and P2, capturing activity over central and centro-parietal midline and adjacent sites. The occipital ROI comprised O1, Oz, O2, PO3, POz, and PO4. Frontal ROIs were divided into left (F5, F3, F1, FC5, FC3, FC1, AF3, F7, FT7) and right (F2, F4, F6, FC2, FC4, FC6, AF4, F8, FT8) regions, spanning dorsolateral, fronto-central, and anterior frontal areas. Temporal ROIs were similarly divided, with the left temporal ROI including T7, C5, TP7, CP5, P7, and P5, and the right temporal ROI including T8, C6, TP8, CP6, P8, and P6, covering superior, middle, and inferior temporal sites as well as temporo-parietal regions. These latter ROIs were retained for analyses within the NH group to provide a more complete characterization of audiovisual prediction under optimal (normal-hearing) conditions. However, temporal ROIs were not included in CI–NH contrasts because recording from temporo-parietal sites was limited by the presence of the cochlear implant, as described above.

To reduce analytical complexity and guide the selection of frequency bands and time windows, we first computed a grand-average time-frequency representation by pooling data across all conditions, participants, and ROIs. This data-driven summary (i.e., derived from the global pattern of power changes rather than predefined boundaries) revealed clear regions of stronger power modulation, from which we defined four frequency bands (8-11 Hz, 12-15 Hz, 16-20 Hz, and 21-30 Hz) and two time windows (0-0.4 s and 0.4-0.8 s relative to gap onset) corresponding to the most prominent modulations (see Supplementary Figure S2). For statistical analysis, we implemented linear mixed-effects models on participant-level averaged data. We used sum contrast coding, so that the intercept reflects the grand mean and each coefficient indexes how much a given level (or level combination, for interactions) deviates from this overall mean, allowing direct tests of main and interaction effects against the grand average.

To make the most of our limited sample size, we adopted a two-step strategy in which NH participants were examined first as a reference group. Because the study is exploratory and the design combines multiple experimental factors, a fully confirmatory, all-in-one search across the entire space of effects would be statistically inefficient and difficult to interpret with a small cohort. We therefore used the NH group to identify where, in a typical processing context, power modulations plausibly indexing predictive mechanisms emerged, thus identifying the most informative time window(s), frequency range(s), and scalp ROI(s). This step served to constrain the subsequent analyses, reducing the dimensionality of the problem and limiting the number of post hoc comparisons. Having established the candidate spectro-temporal-spatial features in NH listeners, we then focused on the CI group to test whether the same signatures were present or altered, thus observing the possible effects of chronically degraded auditory input. In short, prioritizing NH analyses provided an empirical anchor and the flexibility required by the complexity of the design, while preserving sensitivity for detecting group-specific deviations in CI users in the context of a small, hard-to-recruit sample.

Consequently, in analyzing NH participants, the predictors consisted of Constraint (HC/LC), MouthVisibility (V+/V-), FrequencyBand (8-11, 12-15, 16-20, 21-30 Hz), and ROI (frontal left, frontal right, central, temporo-parietal left, temporo-parietal right, occipital) in full interaction (i.e., *Constraint * MouthVisibility * FrequencyBand * ROI*), with Subject modeled as a random intercept and random slope of Constraint by Subject (i.e., *(1 + Constraint | Subject)*). The two temporal windows (0-0.4 s and 0.4-0.8 s) were analyzed separately to focus on temporally local modulations and avoid overly complex models. Interactions were further explored using marginal estimated means computed with the *emmeans* package (Lenth, 2021), allowing pairwise comparisons while controlling for the repeated-measures structure of the data. Subsequently, CI users were analyzed, with predictors selected based on the effects observed in NH controls. This approach ensured that the analysis focused on the relevant time-window, frequency bands and/or ROIs identified in the control group, optimizing sensitivity despite the limited sample size. As additional exploratory analyses, we also modeled the CI group separately to highlight potential group-specific effects.

#### 2.6.3 Event-related potentials

The component of interest in the present study was the N400. Analyses were therefore focused on the average activity across channels in the central ROI, typical of the N400. The time window for averaging was determined by visual inspection of the grand-averaged ERPs. In the ERPs time-locked to target onset, the latency of the N400 modulation appeared slightly delayed relative to the typical 300-500 ms post-target (with onset as early as 200 ms in auditory paradigms; Kutas & Van Petten, 1994), emerging instead starting from around 500 ms after target onset and extending until 1000 ms and even beyond in NH. We discuss this aspect in the Discussion section below (§ 4.3). A linear mixed-effects model was fitted to subject-averaged data, with the categorical predictors (Group, Constraint, MouthVisibility) coded using sum contrasts, with random intercepts for participant and a random slope for Constraint by participant (R formula: *mean_eeg ∼ Group * Constraint * MouthVisibility + (1 + Constraint | ID)*). Given that linear mixed-effects models can effectively handle missing data and unequal group sizes (Maas & Hox, 2005; Pinheiro & Bates, 2000), we retained all participants in the NH group despite the exclusion of one participant from the CI group during ERP statistical analyses.

To further assess robustness, as supplementary analyses we applied these pipelines to ERP data time-locked to gap onset and voice onset (the latter determined by adding trial-level latency of voice onsets measured manually by using Praat; Boersma, 2001). Different event-locking implies different baselines; this approach enables assessment of both the consistency of possible effects and the impact of different baseline choices, allowing for a more informed and nuanced interpretation of the results.

#### 2.6.4 Correlations between pre-target power and language scores

We examined whether interindividual variability in power modulations within the previously identified spectro-temporal window scales continuously with linguistic skills, irrespective of whether a categorical effect reaches statistical significance within each group. In other words, whereas group-wise analyses probe the presence or absence of mean differences, correlational analyses capture graded, cross-participant relationships that can cut across group boundaries and reveal a dimension of variability that is not evident from group contrasts alone. First, we conducted Principal Component Analysis (PCA) on the language comprehension (RTs and accuracy from the lexical decision for word items and the SPM) and production (word counts from semantic and phonological fluency and the sentence generation score) tasks separately, obtaining composite scores that consistently captured performance in each task set. To this aim, we first reverse-coded reaction times (RTs transformed into –RTs), so that higher values (less negative RTs and more positive accuracies) uniformly reflect better performance and positive PC loadings can be interpreted unambiguously as more proficient behavior. We then selected the first component of each task set and identified possible outliers in language scores PCA scores using the 1.5×IQR criterion. Any outliers were removed before performing correlations between individual composite scores and power modulations (specifically, power differences between HC and LC conditions in the 12-15 Hz range, second half of the gap, V+ condition, in the frontal left and central ROIs). To assess the stability of EEG–behavior correlations, we implemented a multi-metric robustness pipeline. For each correlation pair, we computed Pearson and Spearman coefficients and evaluated robustness through: (i) non-parametric bootstrap confidence intervals (2000 resamples; robust if the 95% CI did not include zero; Zou, 2007), (ii) leave-one-out stability (all leave-one-out correlations retaining the sign of the full estimate and SD < 0.05; de Rooij & Weeda, 2020), (iii) method agreement to assess robustness to distributional assumptions and outliers (|Pearson–Spearman| < 0.05), (iv) influential observation diagnostics using Cook’s distance (Cook, 1977), excluding values greater than 4/n, (v) a two-sided permutation test (5000 permutations; robust if the permutation p-value was < .05), and (vi) split-half resampling (2000 random splits; robust if split-half correlations were consistently sign-concordant with the full-sample estimate and the central 95% of split estimates did not include zero). A composite robustness score (0-6) summarized how many criteria were met and was categorized as Low (0-2), Moderate (3-4), or High (≥5) robustness.

## 3. Results

### 3.1 Comprehension tasks: lexical decision and sentence-picture matching

For the lexical decision task (Figure 3A-B), separately for accuracy and RTs, we first compared two generalized linear mixed-effects models with identical fixed effects (type × group), but different random structures. Adding a random slope of type by participant (ID) to the simpler structure with just random intercepts for type and participant significantly improved model fit for both accuracy and RTs (χ²(2) = 86.26, p < 0.001; ΔAIC = 82 for accuracy, and χ²(2) = 423.97, p < 0.001; ΔAIC = 420 for RTs). In other words, the magnitude of the stimulus type effect varied considerably across participants. Accounting for this heterogeneity produced a markedly better-fitting model. Therefore, all reported effects are based on the model with random intercepts for ID and stimulus and a random slope of type for ID (R formula: *accuracy/rt ∼ group * type + (1 + type | ID) + (1 | stimulus*)). The model showed a robust main effect of stimulus type (β = –1.65, SE = 0.32, z = –5.08, p < 0.001), with non-words yielding markedly lower accuracies than words. The group effect was numerically marginal but not significant (β = –0.38, SE = 0.20, z = –1.92, p = 0.055). However, a significant type × group interaction emerged (β = 0.49, SE = 0.20, z = 2.46, p = 0.014), and subsequent contrasts of estimated marginal means revealed that while for non-words CI and NH performed similarly (odds ratio = 1.26, z = 0.44, p = 0.663; CI: 83.5%, NH: 83.9%), for words, CI showed significantly lower accuracy than NH (odds ratio = 0.17, z = –2.90, p = 0.004; CI: 93.8%, NH: 98.1%). As for RTs, the model revealed no reliable group differences, with overall similar latencies for CI users and NH controls (β = 0.10, SE = 0.07, z = 1.38, p = 0.169), although the positive coefficient indicates a tendency for CI users to respond more slowly than NH listeners. In contrast, a robust effect of lexicality (stimulus type), with responses to non-words substantially slower than to words (β = 0.34, SE = 0.05, z = 7.06, p < 0.001), corresponding to roughly a doubling of RTs for non-words relative to words. The Group × Type interaction was small and non-significant (β = −0.02, SE = 0.04, z = −0.45, p = 0.653), indicating that the lexicality effect on RTs was comparable in CI and NH participants.

**Figure 3.**
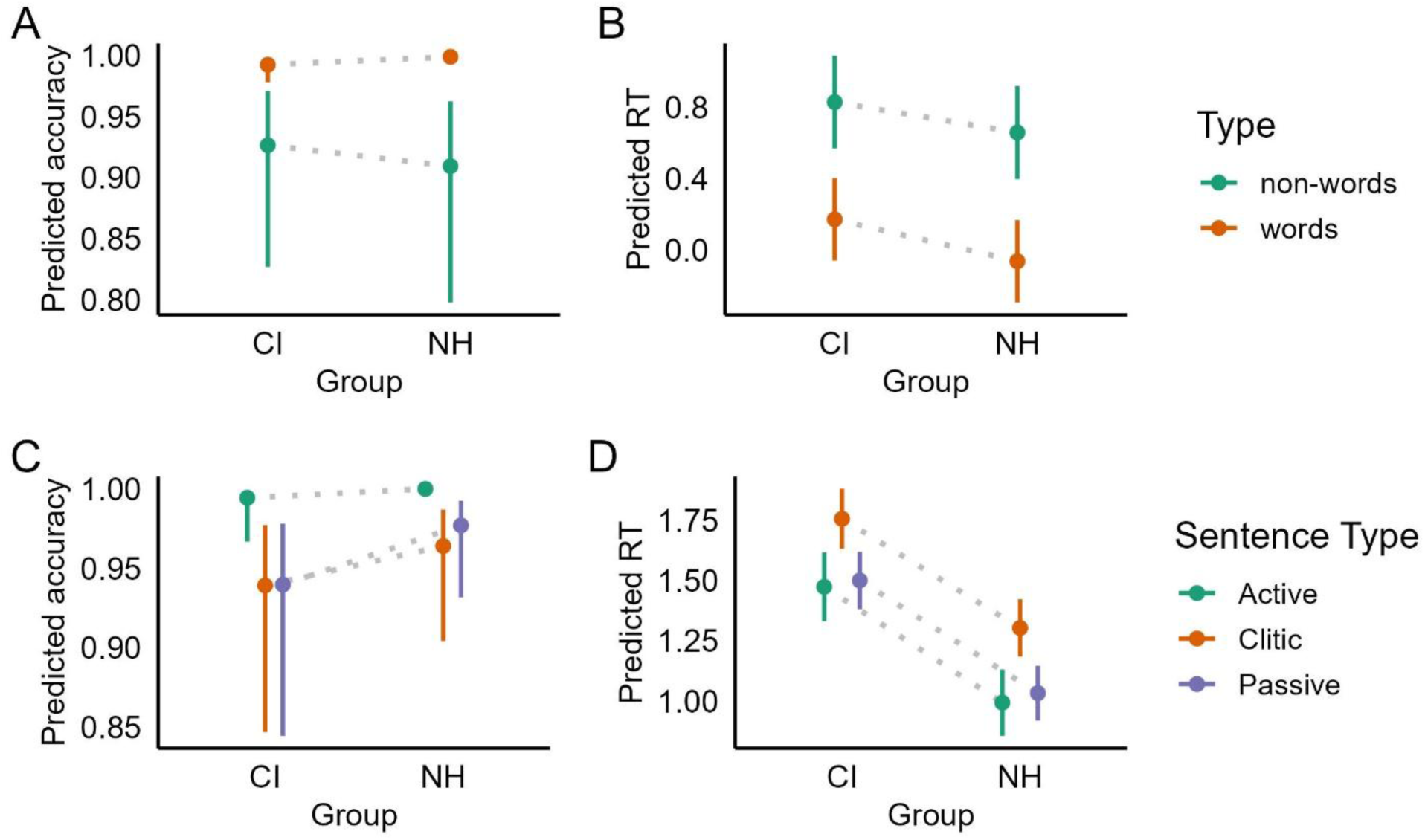
Comprehension tasks. A) Lexical decision: accuracy. B: Lexical decision: RTs. C) Sentence-picture matching: accuracy. D) Sentence-picture matching: RTs. For each measure, estimated marginal means are displayed, along with 95% confidence intervals. Note: For RTs In the SPM task (panel C), confidence intervals are not displayed for NH-Active (ceiling effect: 100% accuracy, 0 errors in 144 trials) due to complete separation in the binomial model, which produces unbounded confidence intervals (0-1) that are uninformative and thus are omitted for visual clarity.

For the SPM task (Figure 3C-D), we proceeded in the same way. We first assessed model fit with simpler and more complex random structures. When modeling accuracy, adding a random slope introduces convergence issues. Even by relaxing the optimization parameters, the model including random slope was not a better fit (χ²(5) = 7.37, p = 0.19; ΔAIC = 2.62). Therefore, for SPM accuracy, a simpler model was fitted, with only random intercepts for participant and item. In the accuracy analysis, several conditions, most notably the NH Active condition, which showed 100% accuracy, displayed performance at or near ceiling, resulting in quasi-complete separation in the logistic mixed-effects model (Clark et al., 2023). This produced extremely large and unstable GLMM coefficients with inflated standard errors (e.g., β for type1 = 6.73, SE = 45.98; β for group1 = −2.94, SE = 22.99), making the logit-scale estimates difficult to interpret. In contrast, the estimated marginal means (EMM) provided stable, meaningful predicted probabilities that closely matched the empirical data: CI users achieved 97.2% accuracy for Active sentences, 87.2% for Clitic, and 85.8% for Passive sentences, while NH listeners performed at ceiling for Active (100%) and near ceiling for Clitic (92.0%) and Passive (91.3%). Pairwise comparisons of estimated marginal means within groups showed significant differences only for CI users, who were more accurate on Active than on Clitic or Passive sentences (OR = 11; p = 0.03), whereas no type-related contrasts were reliable within the NH group due to ceiling performance. Between-group comparisons conducted within each sentence type likewise revealed no significant CI-NH differences, with odds ratios close to 1 and non-significant z-tests (all p > 0.069). Together, these results underscore that EMMs offer a more interpretable and numerically stable summary of accuracy patterns than the raw GLMM coefficients in contexts where ceiling effects induce complete or quasi-complete separation. For RTs, adding the random slope for type by participant improved model fit significantly (χ²(5) = 37.82, p < 0.001; ΔAIC = 28). The more complex model showed significantly slower values in CI users than in NH listeners (β = 0.23, SE = 0.04, p < 0.001). Both sentence-type contrasts were significant: responses were faster to Active than to the other structures (type1: β = –0.11, SE = 0.035, p = 0.002) and slower to Clitic than to Passive/Active sentences (type2: β = 0.19, SE = 0.033, p < 0.001). Pairwise comparisons within each group by computing estimated marginal means confirmed this pattern: both CI and NH participants responded significantly slower to Clitic than to Active sentences (CI: estimate = –0.28, SE = 0.07, p < 0.001; NH: estimate = –0.31, SE = 0.068, p < 0.001) and significantly slower to Clitic than to Passive sentences (CI: estimate = 0.26, SE = 0.066, p < 0.001; NH: estimate = 0.27, SE = 0.062, p < 0.001), while Active-Passive differences were non-significant in both groups (all ps > 0.8). No interactions with group were found, indicating that the syntactic RT pattern was comparable across CI and NH listeners.

### 3.2 Production tasks: verbal fluency and sentence generation

Across tasks, the CI group showed consistently lower scores than the NH group: semantic fluency was significantly reduced in CI participants (M = 16.8, SD = 4.16) compared to NH (M = 19.8, SD = 3.90), U = 95.5, p = 0.036, r = 0.35 (moderate). Sentence generation showed a similar pattern, with CI scoring lower (M = 10.1, SD = 3.31) than NH (M = 12.6, SD = 1.38), U = 84.5, p = 0.014, r = 0.41 (moderate). In contrast, phonological fluency differences were smaller and non-significant (CI: M = 10.7, SD = 5.42; NH: M = 12.4, SD = 4.53), U = 118, p = 0.173, r = 0.23 (small).

### 3.3 Audiovisual sentence comprehension

Comprehension accuracy in the audiovisual sentence comprehension task was high in both groups, with slightly higher performance in the NH group (98.9%, SD = 2.57) than in the CI group (96.3%, SD = 6.12). Only one participant, belonging to the CI group, showed accuracy below 90% (CI_04: 73.8%). Comprehension responses were not further analyzed, as they were only introduced to make sure participants attended the stimuli.

### 3.4 Time-frequency representations

The analysis in the first half of the gap (0-0.4 s) in the NH group, the variables of interest (Constraint and MouthVisibility) did not show reliable effects on power: neither their main effects nor their interaction reached significance (all |t| ≤ 0.93, p ≥ 0.36), and no higher-order interactions involving Constraint × MouthVisibility were significant. In contrast, there were robust main effects and interactions involving ROI and FrequencyBand, which reflect expected physiological differences in the spatial and spectral distribution of endogenous power rather than theoretically relevant modulation by our experimental manipulations.

In the second half of the gap (0.4-0.8 s), there was no reliable main effect of Constraint (β = –0.70, SE = 0.60, t = –1.15, p = 0.26) or Mouth Visibility (β = 0.07, SE = 0.39, t = 0.17, p = 0.86). The Constraint × Mouth Visibility interaction also failed to reach conventional significance (β = –0.64, SE = 0.39, t = –1.63, p = 0.10), although descriptively the constraint effect tended to be more negative when the mouth was visible than when it was occluded. However, importantly, we found a three-way interaction Constraint × MouthVisibility × FrequencyBand (β = –1.63, SE = 0.68, t = –2.39, p = 0.017). Because we used sum-to-zero contrasts, the significant interaction term indicates that the Constraint × MouthVisibility interaction deviates from its grand-mean pattern specifically in the 12-15 Hz band relative to the 8–11 Hz band, averaged over ROIs: the negative coefficient indicates that in 12-15 Hz the HC-LC difference between V+ and V-is less positive / more negative than in 8-11 Hz, whereas no effect was detected for the other bands (Figure 5A-B). Post-hoc estimated marginal means confirmed that the only significant HC–LC contrast emerged in the V+ condition within the 12–15 Hz band (estimate = –5.76, SE = 2.41, t = –2.39, p = 0.018). All other MouthVisibility × FrequencyBand combinations showed no reliable differences (all |t| ≤ 1.382, all p ≥ 0.168; Figure 5C).

**Figure 4.**
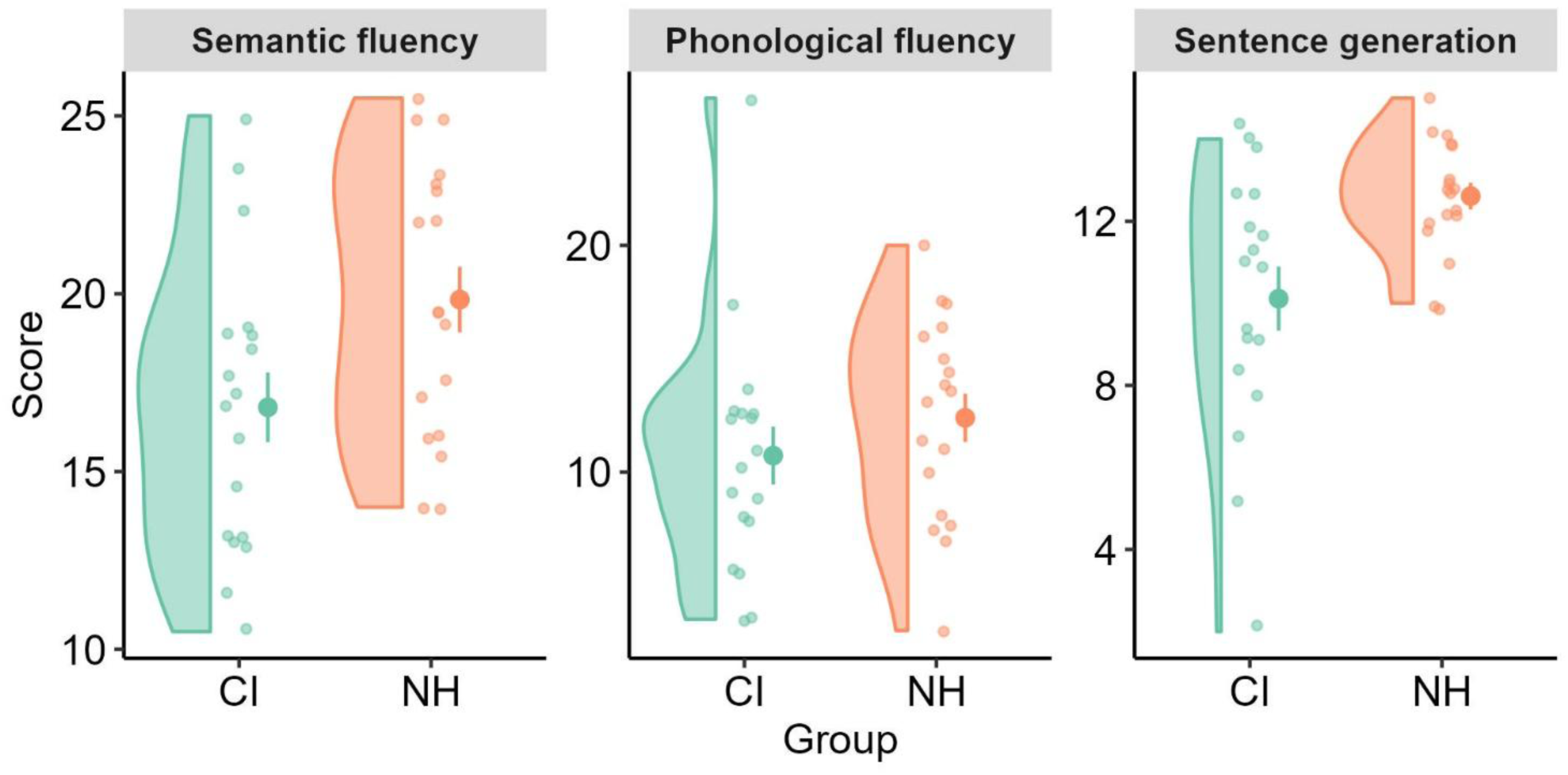
Production tasks. For each measure, individual raw scores, distributions, means and standard errors are shown. Note that the difference in the phonological fluency task is not significant even when excluding the CI outlier with score > 25 (mean = 9.79, SD = 3.83, U = 100, p = 0.085, r = 0.294).

**Figure 5.**
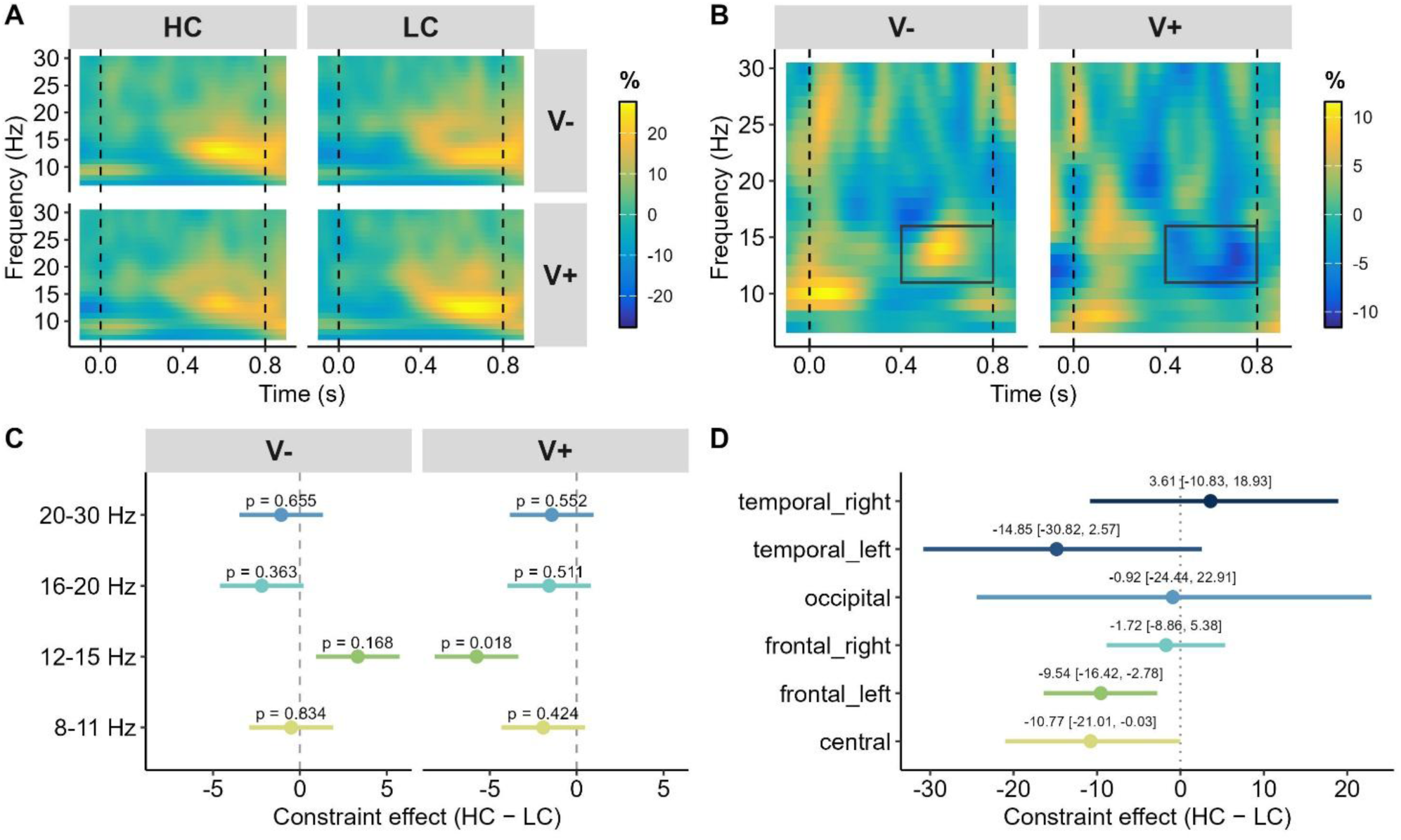
Constraint effect in normal hearers (NH). A) Time-frequency representations during the silent gap for HC and LC contexts, separately for mouth visibility (V-= covered; V+ = visible). Gap onset (0) and offset (0.8) are marked with dashed vertical lines. Color indicates %-power change relative to baseline (pre-sentence interval). B) Constraint effect (HC-LC) in the two mouth visibility condition averaged across all ROIs (main effect of Constraint), with a rectangle highlighting the time-frequency window where the difference is statistically significant. Gap onset (0) and offset (0.8) are marked with dashed vertical lines. Color indicates %-power change relative to baseline (pre-sentence interval). C) Estimates marginal means of the constraint effect (HC-LC) for each frequency range. D) Bootstrapping analysis showing mean and 95%CI range of the constraint effect across the ROIs (12-15 Hz, V+ condition).

To exploratory assess the spatial distribution of the constraint effect in the NH group, we ran a non-parametric bootstrap analysis on time-frequency power. We first restricted the data to NH participants, the 12-15 Hz band, visible-mouth trials (V+), and the 0.4-0.8 s pre-target interval. For each Subject × Constraint (HC, LC) × ROI combination, we computed the mean power across time points, yielding one value per subject, condition, and ROI. Then, separately for each ROI, we used a bootstrap procedure with 1000 resamples (*boot* package v1.3 in R, Canty & Ripley, 2024). The subject-level observations were resampled with replacement and, at each iteration, recomputed the mean power for HC and LC and their difference (HC-LC). The resulting bootstrap distribution of HC-LC differences was used to derive (i) the bootstrap estimate of the effect (mean of the distribution), (ii) 95% percentile confidence intervals, and (iii) a two-tailed empirical p-value obtained as twice the smaller of the proportions of bootstrap estimates above and below zero (computed as 2 × min[P(difference < 0), P(difference > 0)]). This procedure provides a robust, distribution-free estimate of the spatial pattern of the Constraint effect without assuming normality or homogeneity of variance.

The analysis revealed a clear spatial pattern (Figure 5D). The constraint effect (HC − LC) was reliably negative over the left frontal and central ROIs, indicating lower 12–15 Hz power for HC than LC sentences in these regions. In the left frontal ROI, the mean effect was –9.54 (95% CI [-16.42, –2.78], p = 0.008), and in the central ROI it was –10.77 (95% CI [-21, –0.03], p = 0.048). A similar, but statistically not significant, trend was observed over the left temporal ROI (–14.85, 95% CI [-30.82, 02.57], p = 0.068). In contrast, no effects were found in the right frontal (–1.72, 95% CI [-8.86, 5.38], p = 0.658), occipital (–0.92, 95% CI [-24.44, 22.91], p = 0.918), and right temporal ROIs (3.61, 95% CI [-10.83, 18.93], p = 0.586). Overall, the results suggest that the constraint-related reduction in upper-alpha/low-beta power is primarily left-lateralized and maximal over frontal and central regions, with a less stable contribution from the temporo-parietal region.

Having identified the relevant frequency band, ROIs, time window, and mouth-visibility condition in NH participants in the first step, we then restricted the CI analyses to this spectro-temporal window in the V+ condition. This targeted approach allowed us to assess whether the same signature observed in normal hearing conditions was preserved in CI users or instead altered under chronically degraded auditory input. In CI users, this effect was not found; instead, the estimate showed a statistically non-significant numerical trend in the opposite direction (β = 0.631, SE = 1.334, t = 0.473, p = 0.643; see Figure 6A-C).

**Figure 6.**
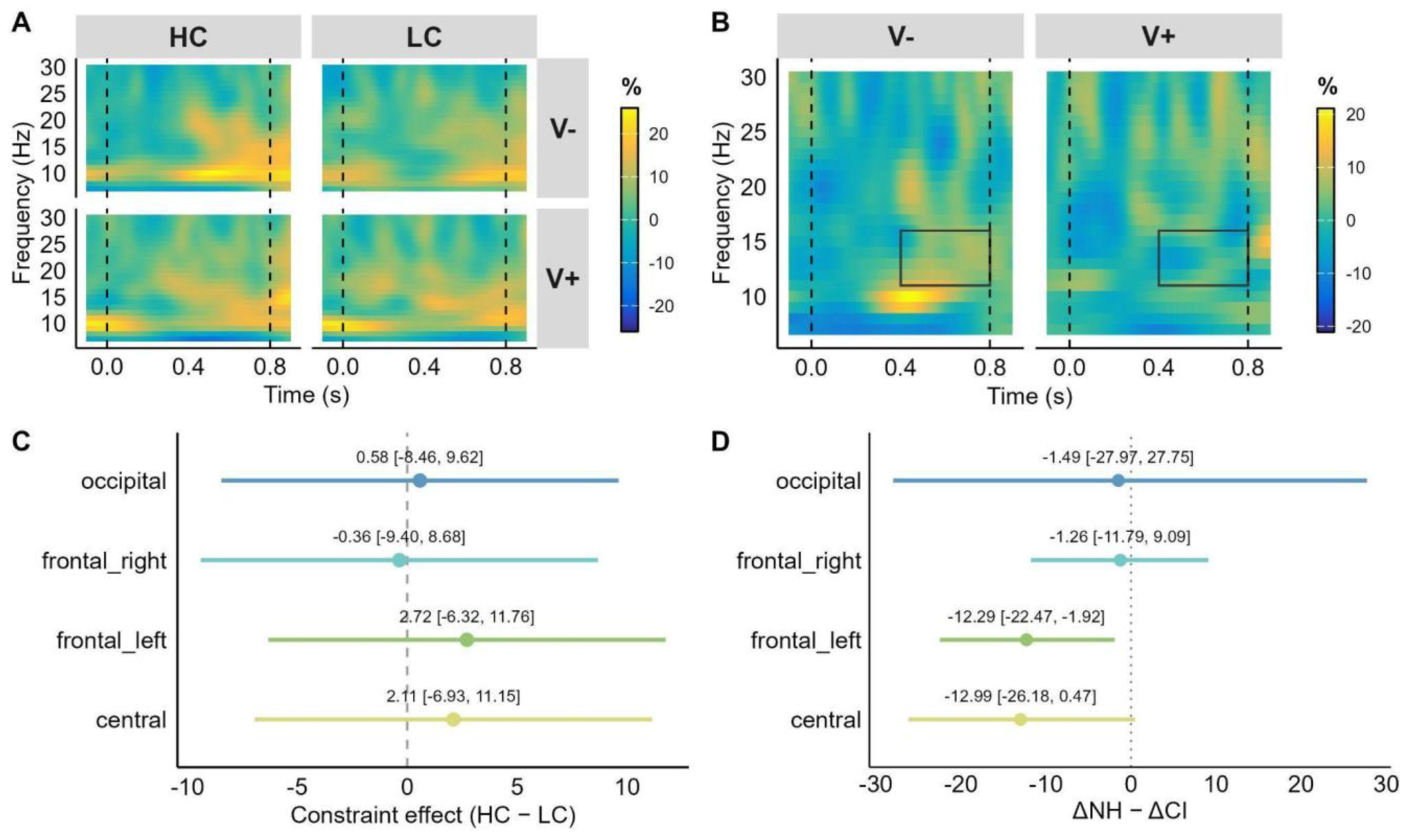
Constraint effect in cochlear implant users (CI). A) Time-frequency representations during the silent gap for HC and LC contexts, separately for mouth visibility (V-= covered; V+ = visible). Gap onset (0) and offset (0.8) are marked with dashed vertical lines. Color indicates %-power change relative to baseline (pre-sentence interval). Temporo-parietal sensors are excluded. B) Constraint effect (HC-LC) in the two mouth-visibility condition averaged across ROIs, with a rectangle highlighting the time-frequency window where the effect was detected in the NH group and where the analysis is focused. Gap onset (0) and offset (0.8) are marked with dashed vertical lines. Color indicates %-power change relative to baseline (pre-sentence interval). Temporo-parietal sensors are excluded. C) Estimates marginal means of the constraint effect (HC-LC) for each ROI, showing no power decrease effect in any sensor cluster in the V+ condition in the time-frequency window of interest. D) Bootstrapping analysis of the constraint effect (12-15 Hz, V+ condition) between CI and NH showing mean and 95%CI range.

To further restrict the spatial specificity of the group differences, we performed bootstrapping analysis of the Constraint difference between the groups. Mean power was computed restricted to V+ and 12-15 Hz, separately for each Subject, Group, Constraint, and ROI (excluding temporal sites for comparison between the groups). For each ROI, we quantified the constraint difference as HC – LC within each group and then calculated the group difference, defined as ΔGroup = (HC – LC)_NH_ − (HC – LC)_CI_. To evaluate the reliability of this group difference without relying on parametric assumptions, we used a nonparametric bootstrap procedure with 1000 resamples. On each bootstrap iteration, subjects were sampled with replacement within each group, preserving the original data structure (i.e., each selected subject contributed their full set of HC and LC values for that ROI). For the resampled dataset, we recomputed mean HC and LC power within each group, derived the contrast HC – LC for NH and CI, and calculated a new estimate of ΔGroup. This yielded an empirical bootstrap distribution of 2000 ΔGroup values for each ROI. From this distribution, we extracted percentile-based 95% confidence intervals, reflecting the range of ΔGroup values compatible with the observed data. Two-tailed bootstrap p-values were computed as twice the proportion of bootstrap samples with a sign opposite to the observed effect. This approach provides a direct, distribution-free estimate of the stability and direction of the group difference in constraint-related low beta power modulation. By applying this procedure independently to each ROI, we identified the group of sensors that contributed most reliably to the divergence between NH and CI listeners in predictive pre-target beta activity. The bootstrap analysis (Figure 6D) showed that the group difference (NH – CI) in the Constraint effect was most pronounced over the left frontal region (mean = –12.29, 95% CI [-22.47, –1.92], p = 0.026), with a similar but slightly less reliable trend over the central ROI (mean = –12.99, CI [-26.18, 0.47], p = 0.058). In contrast, effects were negligible and unreliable over right frontal (mean = –1.26, CI [-11.79, 9.09], p = 0.848) and occipital regions (mean = –1.49, CI [-27.97, 27.75], p = 0.898). Taken together, the findings indicate that the group difference is more likely to reflect a spatially circumscribed effect, primarily involving left frontal sensors and, less robustly, central sensors. However, the lack of data in temporo-parietal sensors in the CI group prevents us from assessing whether and how the neural dynamics between groups differ in those ROIs.

We also analyzed the CI group independently with a full model as initially for NH. During the first half of the gap, the effect of Constraint was non-significant (β = –0.12, SE = 0.71, t = –0.17, p = 0.87). By contrast, Mouth Visibility showed a reliable main effect (β = 1.22, SE = 0.51, t = 2.41, p = 0.016), corresponding to higher values when the mouth was visible than when it was occluded. However, this visibility effect was not further modulated by Constraint (β = 0.02, SE = 0.51, t = 0.04, p = 0.97). Likewise, all higher-order interactions involving Constraint and/or Mouth Visibility with Frequency Band and ROI (two-, three-, and four-way terms) were small in magnitude and nonsignificant (all |t| ≤ 1.16, all p ≥ 0.25), indicating that neither the presence of a semantic constraint nor mouth occlusion systematically shaped the spatio-spectral pattern of the effect. In the second half of the gap, while the effect of Constraint did not reach conventional significance threshold (β = 1.273, SE = 0.624, t = 2.039, p = 0.057), the numerical trend was in the opposite direction of the expected one (i.e., slightly higher power for HC than for LC condition), in line with the previous analysis. The effect of Mouth Visibility (β = –0.10, SE = 0.50, t = –0.20, p = 0.84) and the Constraint × Mouth Visibility interaction (β = 0.04, SE = 0.50, t = 0.07, p = 0.94) were null, indicating that mouth occlusion did not significantly alter pre-target power modulations, and that any effect of Constraint did not differ between visible and occluded mouth conditions. Likewise, all higher-order interactions involving Constraint and/or Mouth Visibility with Frequency Band and ROI (two-, three-, and four-way terms) were small in magnitude and nonsignificant (all |t| < 1.52, all p ≥ 0.13), suggesting that the pattern of results across frequency bands and ROIs did not depend reliably on semantic constraint or mouth visibility.

### 3.5 Event-related potentials

The analyses showed a significant main effect of Group, with a negative coefficient indicating more positive mean EEG in NH than CI (β = –1.13, SE = 0.34, t = –3.29, p = 0.002). There was also a strong main effect of Constraint, such that LC sentences elicited more negative amplitudes than HC ones (β = 0.92, SE = 0.12, t = 7.52, p < 0.001). The main effect of MouthVisibility showed only a numerical trend, with a negative coefficient meaning slightly more positive amplitudes when the mouth was visible (V+) than when it was occluded (V-; β = –0.21, SE = 0.12, t = –1.68, p = 0.096). None of the two-way interactions reached significance: Group × Constraint (β = –0.08, SE = 0.12, t = –0.69, p = 0.49), Group × MouthVisibility (β = 0.20, SE = 0.12, t = 1.60, p = 0.11), and Constraint × MouthVisibility (β = 0.12, SE = 0.12, t = 1.01, p = 0.32). However, there was a significant three-way interaction Group × Constraint × MouthVisibility (β = –0.30, SE = 0.12, t = – 2.41, p = 0.018), indicating that the way the semantic constraint effect (HC vs LC) depends on mouth visibility (V-vs V+) differs between CI users and NH listeners. More specifically, the modulation of mouth visibility during the previous sentence frame affected LC (non-predictable) targets only in the NH group, with more negative values for V-relative to V+ (see Figure 7).

**Figure 7.**
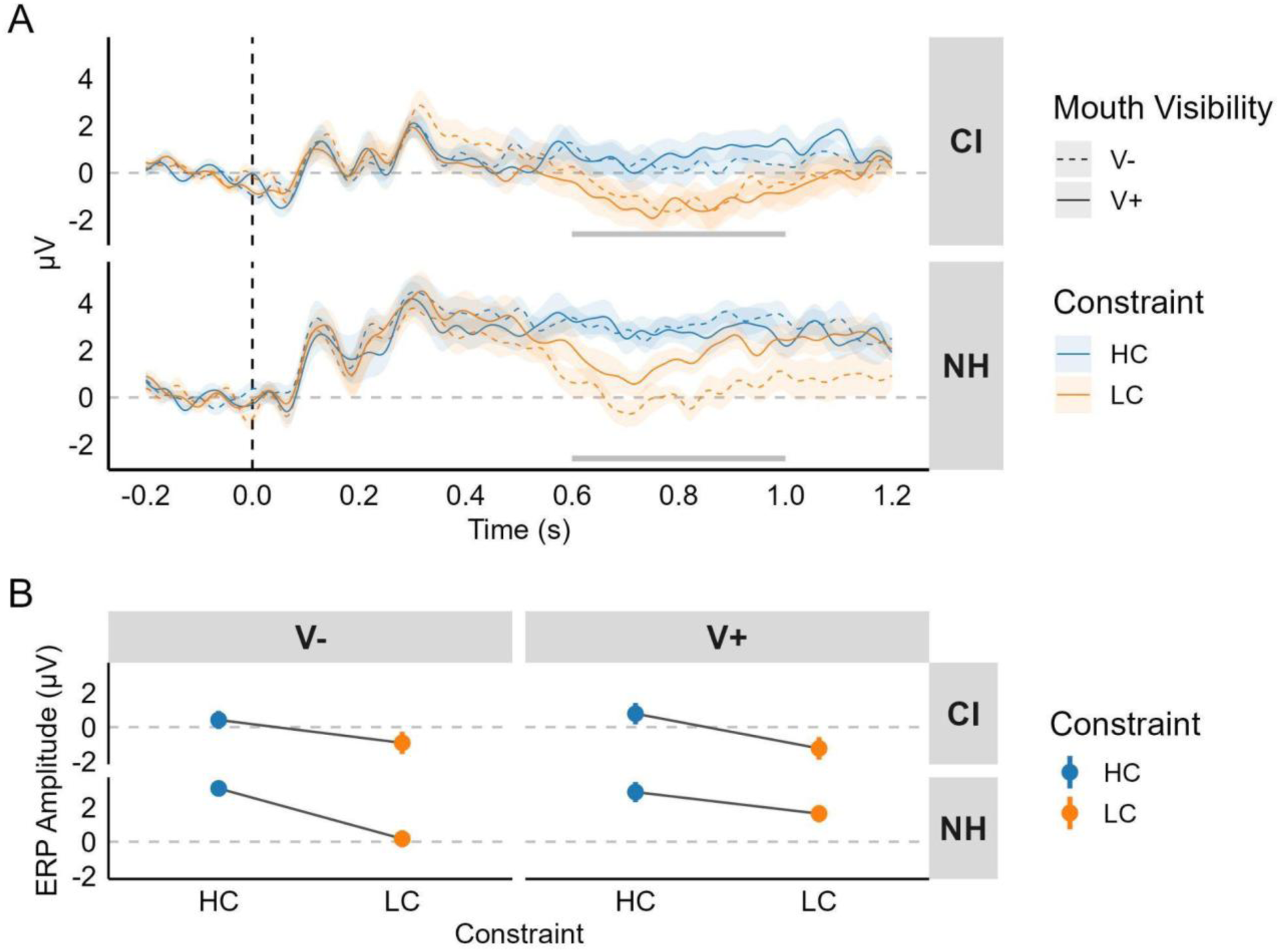
Target-locked ERP. Panel A shows the time-series of the ERP separately for group. Time = 0 represents the onset of the audio-video file of the target word. The time-window highlighted with the grey line represents the interval where statistical testing has been performed. The boxplots show the jitter between target video file actual audio speech and mouth movement onsets, which explains the temporal delay of the N400. Panel B shows means and standard deviations of the voltage in the time-window marked in panel A. The data was averaged across the central ROI defined in the Methods section, which included C1, Cz, C2, CP1, CPz, CP2, P1, Pz, P2.

However, we reasoned that the original baseline choice may have introduced distortions, especially because the –200 to 0 ms pre-target interval likely contains neural activity related to prediction generation. To examine the robustness of the Constraint × Group interaction, we therefore computed ERPs time-locked to the gap onset and applied a pre-gap baseline (–200 to 0 ms), which minimizes the subtraction of activity that may be functionally relevant (see Figure 8). This baseline is suboptimal for the component of interest, as downstream ERP effects are attenuated when the baseline and the component are far apart (Luck, 2014). Indeed, the N400 effect is substantially reduced. Nonetheless, this analysis provides an important check on whether the initial baseline choice might have introduced distortions in the windows of interest.

**Figure 8.**
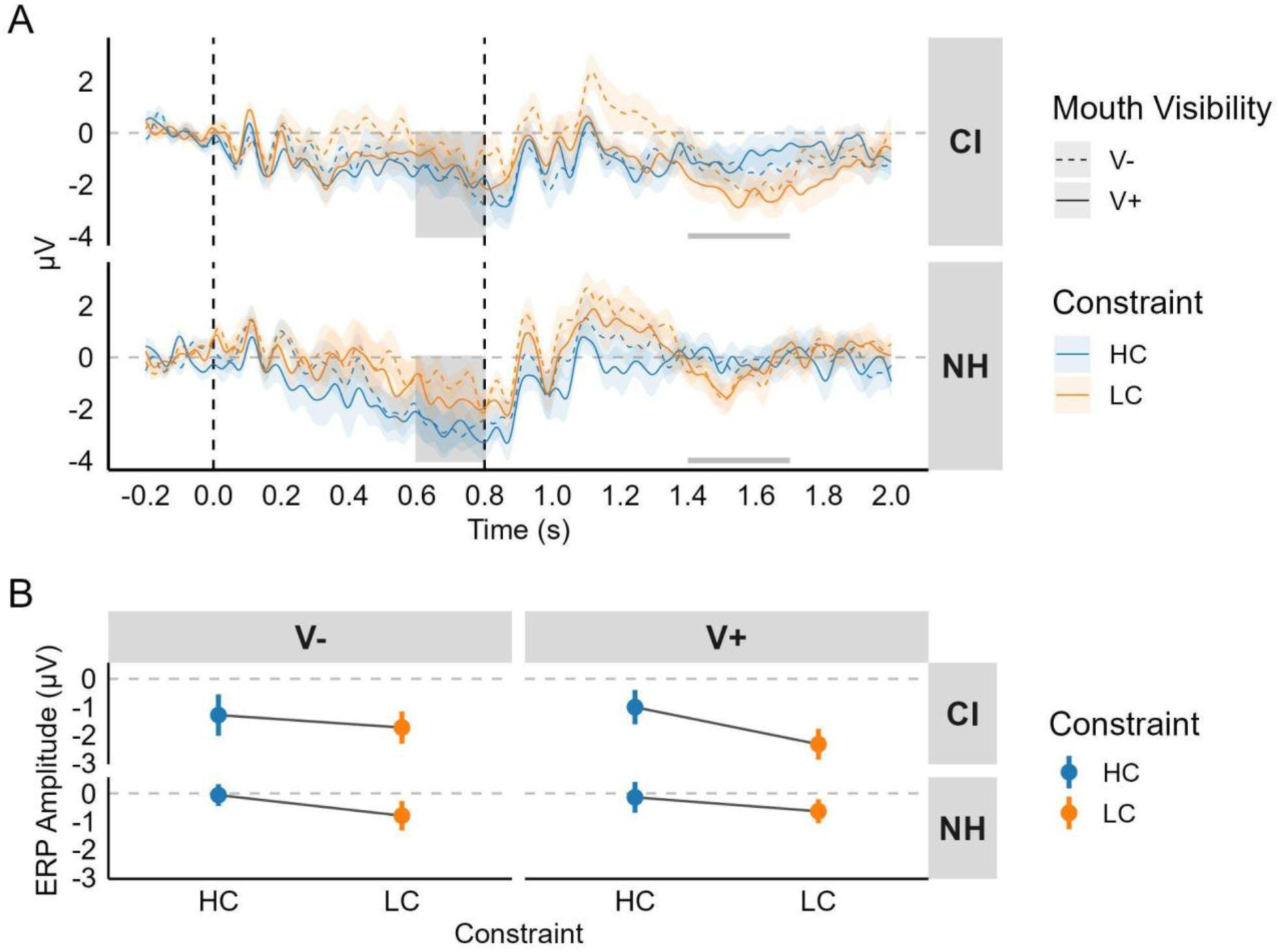
Gap-locked ERP. Panel A shows the time-series of the ERP separately for groups. Time = 0 represents the onset of the silent gap after the sentence frame; time = 0.8 represents the end of the gap. The time-window highlighted with the grey line represents the interval where statistical testing has been performed. Data has been magnified to facilitate visualization. The shaded area between t = 0.6 and t = 0.8 represents the baseline interval that is used to compute target-locked ERPs, showing that subtracting that activity generates the artefactual interaction in the later time window of the target-locked ERP (see Figure 7). Panel B shows means and standard deviations of the voltage in the time-window marked in panel A. The data was averaged across the central ROI defined in the Methods section, which included C1, Cz, C2, CP1, CPz, CP2, P1, Pz, P2.

In the time-window of interest (1.4 to 1.7 s time-locked at gap onset), the linear mixed-effects model (same specifications as for target-locked analysis) revealed a significant main effect of Constraint (β = 0.38, SE = 0.17, t = 2.26, p = 0.031), reflecting more negative amplitudes for targets after LC than HC sentences when averaging across groups and mouth-visibility conditions. The effecs of Group did not reach conventional statistical significance (β = –0.59, SE = 0.3, t = –1.957, p = 0.059) and the effect of MouthVisibility was null (β = 0.03, SE = 0.13, t = 0.23, p = 0.81). None of the interactions reached significance (all |t| ≤ 0.38, all p ≥ 0.28), indicating that the Group difference and the Constraint effect did not reliably vary as a function of mouth visibility in this second analysis.

By looking at the time-series in Figures 7 and 8, it appears clear that the interaction found in the target-locked ERP is an artifact introduced as a result of baseline subtraction in the pre-target interval, and the consequent lowering of the LCV-condition in the NH group. This interpretation is supported by the fact that, by applying the same linear mixed-effects models as for the other ERP analyses in the 0.6-0.8 s interval to gap-locked ERPs (corresponding to the last 200 ms of the silent gap), we found a main effect of Constraint (β = –0.51, SE = 0.17, t = – 3.05, p = 0.004). For this reason, we refrain from analyzing and discussing the interaction in the target-locked ERP any further. The only consistent result across multiple plausible analytical approaches in the present ERP data is the presence of a constraint effect.

As follow-up exploratory analyses in the gap-locked ERP, we analyzed the 1.7-1.8 s window by using the same model specification. This was decided on the basis of visual inspection, suggesting that the temporal extension of the N400 is longer for CI users than for NH controls. In this time window we found a 3-way interaction between Group, Constraint and MouthVisibility (β = –0.24, SE = 0.11, t = –2.23, p = 0.029). To further characterize the interaction, we obtained estimated marginal means from the mixed-effects model, then we first computed the simple effect of Constraint (LC–HC difference in mean amplitude) separately within each Group × MouthVisibility combination. We then contrasted these LC–HC effects between groups, separately for each level of MouthVisibility, yielding tests of whether the size of the constraint effect differed between CI and NH when the mouth was occluded (V-) and when it was visible (V+). When the mouth was occluded (V-), the constraint effect did not differ between CI and NH (β = 0.09, SE = 0.72, t = 0.13, p = 0.9). In contrast, when the mouth was visible (V+), the constraint effect was significantly larger (more negative) in CI than in NH (β = –1.85, SE = 0.72, t = –2.57, p = 0.012). This may indicate that when the mouth was visible in the preceding sentence frame, CI users required longer processing time at the target than NH people, while no difference emerges with mouth occlusion during sentence unfolding.

### 3.6 Correlations

The PCA on language skills measures showed that the first principal component (PC1) accounted for 64.78% of the variance in the production task set and 88.27% in the comprehension task set. In both sets, the measures included contributed comparably to PC1, indicating that it captures the shared variance across tasks and provides a reasonable summary index of overall performance. Notably, the much higher variance explained in comprehension suggests a more strongly unitary skill dimension (i.e., speed-accuracy covarying tightly across measures), whereas production showed a less one-dimensional structure, with additional components still capturing meaningful variance (see Supplementary Figure S6). The outlier detection procedure identified one single outlier (CI_04; see Supplementary Figure S7). All subsequent correlations were performed and assessed by excluding this participant. PC1 scores for the comprehension and production measures were correlated with the HC–LC Constraint effect in the 12–15 Hz range during the 0.4–0.8 s gap interval within the left frontal and central ROIs. These ROIs were selected because they had been identified as plausible generators of the effect in the NH group and of the group differences. Correlation robustness checks revealed that two correlations, power_frontal_left ∼ PCA_prod (r = –0.46, p = 0.006, p_FDR_ = 0.022) and power_central ∼ PCA_prod (r = –0.38, p = 0.024, p_FDR_ = 0.049), show high robustness, with bootstrap CIs not crossing zero and high stability across permutation tests. Correlations with PC1 of comprehension scores do not survive FDR-correction and are also found to be unreliable (power_central ∼ PCA_comp: r = –0.34, p = 0.049, p_FDR_ = 0.065; power_frontal_left ∼ PCA_comp: r = –0.26, p = 0.132, p_FDR_ = 0.132). Overall, more reliable effects emerge for PC1 production, suggesting that stronger power decreases in the constraint effect (i.e., prediction generation) is associated with higher production skills scores (Figure 9).

**Figure 9.**
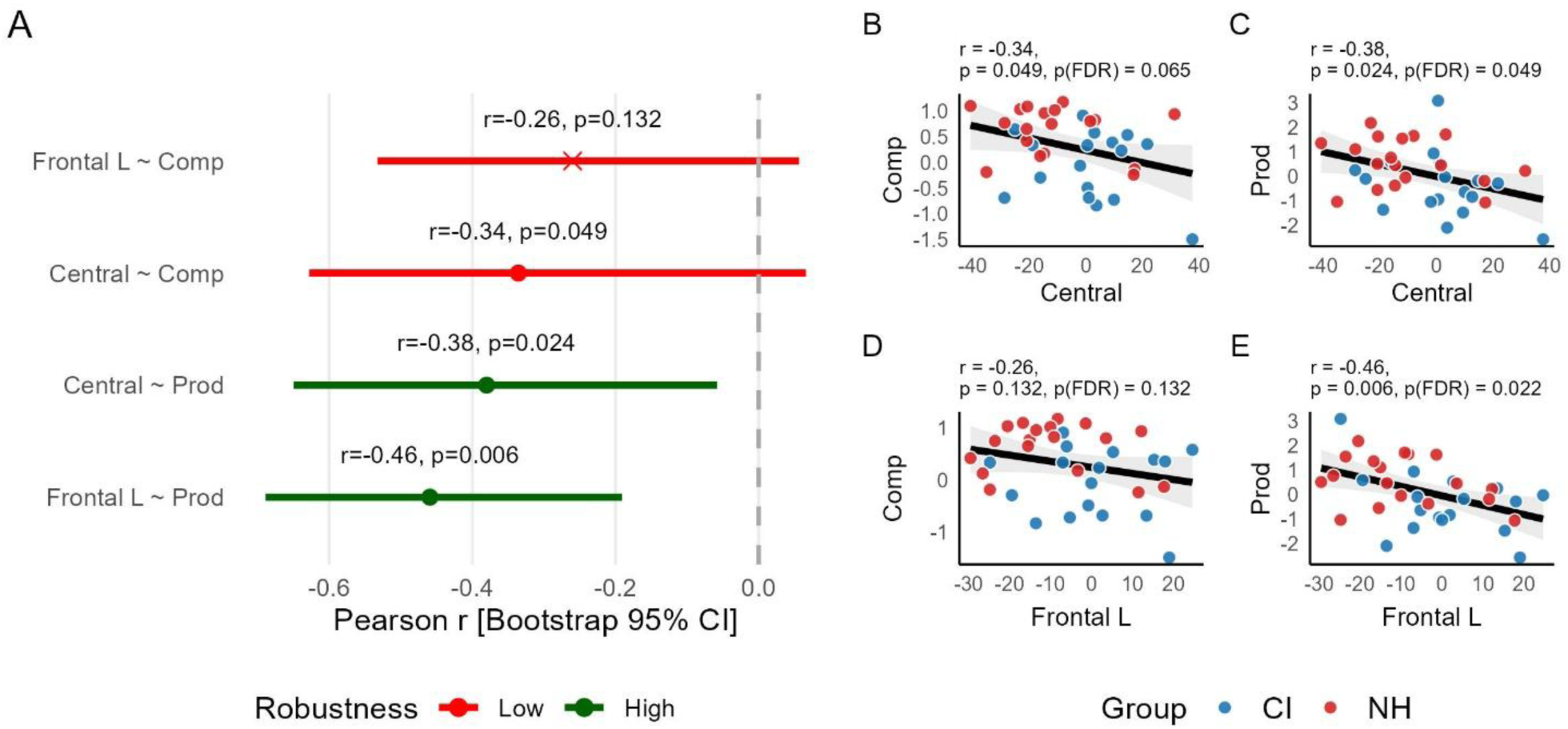
Robustness check on EEG power-language skills correlations. A) Result of the robustness check procedure (non-parametric bootstrap CIs, leave-one-out stability, Pearson-Spearman agreement, and influence diagnostics using Cook’s distance, FDR-correction). The color represents the composite score describing correlation robustness, while the shape represents statistical significance (uncorrected). B-E) Scatterplots for each correlation between PC1 of the production and comprehension scores and power modulations (HC-LC) in the 12-15 Hz of the second half of the gap in the frontal left and central ROIs.

As additional exploratory correlational analysis, to better understand how CI use and audiological measures relate to electrophysiological outcomes, especially for the post-target N400, where no clear group differences emerged with respect to NH controls, we correlated PTA4 scores and years of CI use with the N400 effect (LC–HC) in the two mouth visibility conditions, computed in the target-locked ERP (0.6 – 1 s). The same robustness checks were applied. These analyses showed robust negative correlations between PTA4 and the N400 effect in both visible (r = –0.55, p = 0.022) and occluded mouth (r = –0.53, p = 0.027) conditions (Supplementary Figure S7). More specifically, increasing PTA4 scores (worse auditory threshold) corresponded to bigger N400 effects. Despite FDR-correction making p-values just beyond conventional significance threshold (p_FDR_ = 0.055), the robustness assessments provided high scores of robustness across the methods adopted. No relationships with years of CI use were found. To assess the consistency of these relationships, we did the same analyses with the gap-locked ERP in the extended window 1.4 to 1.9 s post-gap onset. In this analysis, none of the correlations were significant and were all weakly robust (Supplementary Figure S8).

## 4. Discussion

In this study, we compared EEG markers of predictive processing during audiovisual comprehension of high-vs low-constraining sentences in cochlear implant (CI) users and normal hearing (NH) controls, while manipulating mouth visibility (visible, V+; occluded, V-). More specifically, we studied pre-target alpha-beta power and post-target N400 ERP. Based on evidence that articulatory mouth cues are particularly informative in adverse listening conditions, we hypothesized that CI users would show stronger neural markers of prediction generation than NH controls when the mouth was visible, namely, a larger pre-target alpha-beta power decrease and a stronger N400 constraint effect (as the result of a stronger negativity reduction for predicted targets, signaling relative higher ease of processing). This pattern would have indicated greater reliance on visual articulatory information to generate predictions. When the mouth was occluded, we expected to replicate previous findings with auditory-only speech. We further expected NH participants to show overall stronger predictive effects in V-due to their access to a more detailed auditory representation.

The results, however, painted a more complex picture. First, we did not observe the classic alpha-beta power decrease for high-vs low-constraint contexts when the mouth was occluded (V-). Second, when such an effect did emerge (in the V+ condition), it was present only in NH controls, who showed a stronger power decrease in central, left frontal, and left temporo-parietal sensors in the lower beta range (12–15 Hz) during the second half of the gap, whereas CI users did not. Third, the post-target N400 effect was not modulated by group or by mouth visibility, but the effect in V+ seems to last longer in the CI group. Lastly, the lower beta band decrease was negatively correlated with production scores, such that higher production proficiency was associated with more negative power values (i.e., a larger prediction-related power decrease). We will discuss these results in more detail in the next sections.

### 4.1 Covering the mouth with a digital rectangle is not equivalent to an audio-only condition

One unexpected finding was the absence of a constraint effect on pre-target oscillatory power in the mouth-occluded (V-) condition, even in NH controls. This null result was surprising given that numerous studies have reported alpha-beta power decreases in anticipation of predictable targets during purely auditory or written sentence comprehension (Gastaldon et al., 2020, 2023; León-Cabrera et al., 2022; Molinaro & Monsalve, 2018; Rommers et al., 2017; Roos at al., 2024; Wang et al., 2018; but see Huizeling et al., 2023 for a study that does not replicate this typical pattern in another complex visual setting such as visual world paradigm in virtual reality). Several factors may account for this discrepancy.

First, the presence of a visible speaker, even with the mouth covered, likely introduced a degree of audiovisual integration that differs from the strictly auditory-only conditions typically used in studies reporting robust pre-target oscillatory effects. Second, in this study we interleaved V+ and V-conditions rather than presenting them in separate blocks. The unpredictable alternation between mouth-visible and mouth-occluded trials may have induced a sustained audiovisual processing mode that persisted even when the mouth was not visible. Under such conditions, listeners may have adopted a general strategy of attending to and integrating visual information whenever it was available, rather than flexibly switching between unimodal and multimodal processing on a trial-by-trial basis. This interpretation is consistent with models of multisensory attention suggesting that audiovisual integration is strongly shaped by expectations and supramodal attentional sets about stimulus modality (Talsma et al., 2010), rather than solely by the immediate sensory content of each trial. At the same time, only V+ trials provided rich, temporally precise articulatory information that could reliably constrain forthcoming phonological and lexical candidates. We therefore suggest that the pre-target low-beta decrease reflects a specifically speech-driven predictive mechanism that is strongly engaged when detailed mouth movements are visible. When mouth cues are occluded but a speaker is still present, more general audiovisual integration processes may be recruited, which are not captured by this specific spectro-temporal modulation in this time window. Third, the ecological validity of the mouth-occlusion manipulation deserves consideration. In everyday communication, speakers rarely have their mouths artificially obscured while their faces remain otherwise visible. The resulting combination of a naturalistic face with an opaque, static grey rectangle over the mouth is visually unusual and may have created a conflict between participants’ expectations about how a talking face should look and the incoming visual information. Such conflicts are likely to alter processing dynamics in ways that go beyond the simple absence of articulatory cues. The rectangle itself may have attracted attention or introduced uncertainty about upcoming articulations, disrupting the smooth, automatic use of visual speech that typically supports prediction in natural audiovisual communication. Recent work using “virtual masks” (digitally superimposed masks that hide the mouth while preserving the original audio; Fantoni et al., 2024) shows that this kind of artificial occlusion is a very powerful manipulation: it eliminates lip-reading, eliminates neural tracking of lip movements, and reduces early neural tracking of the speech envelope relative to an unmasked face, even though the acoustic signal is identical. At the same time, the same studies were designed primarily to dissociate visual and acoustic obstacles, rather than to fully reproduce everyday face-to-face communication. In our case, the grey rectangle represents an even less ecological visual configuration than a real face mask, because it selectively removes the mouth region while leaving the rest of the face intact and visually salient. We therefore consider it likely that our mouth-occluded condition engaged an atypical audiovisual processing mode, in which listeners may have down-weighted or even distrusted visual cues rather than using them in the way they would with a naturally visible or naturally masked talker. This could help explain why the pre-target low-beta suppression emerged robustly when rich, temporally precise mouth movements were visible, but not when the mouth was covered by a digital patch, despite the presence of a speaker’s face on the screen.

Taken together, these considerations suggest that future studies aiming to isolate auditory speech processing in the context of audiovisual paradigms may need to employ different approaches, such as presenting auditory-only stimuli without any visual component, or using more ecologically valid visual occlusions (e.g., surgical masks) that might be processed more naturally by the visual system.

### 4.2 Constraint-related pre-target lower beta decrease in NH controls, but not in CI users

The central finding of the present study is that NH controls showed a significant constraint effect in lower-beta power (12–15 Hz) during the second half of the pre-target gap, specifically when the mouth was visible, whereas CI users did not exhibit this effect. The spatial distribution of this effect over left frontal, central, and left temporo-parietal regions is consistent with the involvement of left-lateralized language networks in prediction generation (Federmeier, 2007; Kuperberg & Jaeger, 2016). The emergence of this effect specifically in the mouth-visible condition for NH controls suggests that articulatory visual information enhances or enables the neural mechanisms underlying prediction generation. Visual speech cues provide advance information about upcoming phonetic content, including place of articulation and voicing features, that temporally precedes the acoustic signal (Chandrasekaran et al., 2009; van Wassenhove et al., 2005). This temporal lead of visual information may facilitate the activation of lexical candidates before acoustic information becomes available, thereby amplifying the neural signature of predictive processing (Piazza et al., in preparation). The integration of visual and auditory speech information appears to create conditions that are particularly favorable to robust prediction generation, at least in individuals with intact auditory processing. The absence of this constraint-related low beta decrease in CI users is a striking finding that warrants careful interpretation. Several non-mutually exclusive explanations may account for this result.

● **Degraded auditory input undermines robust prediction generation**. The spectrally impoverished signal delivered by cochlear implants may fail to provide sufficient information for building the rich contextual representations that support prediction. The reduced spectral resolution of CI-mediated hearing particularly impacts the perception of phonetic features that are important for distinguishing between lexical candidates (Zeng et al., 2008). Without reliable bottom-up phonetic information, listeners may be unable to narrow down predictions to specific word forms, even when semantic context strongly constrains interpretation. This account is consistent with evidence that CI users show less efficient anticipatory eye movements in visual world paradigms (Blomquist et al., 2021; Nagels et al., 2020) and delayed predictive pupil responses (Winn, 2016).
● **Resources are allocated to effortful listening**. Processing degraded speech requires increased cognitive effort, as evidenced by pupillometric, behavioral, and neuroimaging studies (Peelle, 2018; Pichora-Fuller et al., 2016; Zekveld et al., 2018). If the cognitive resources required for prediction generation overlap with those needed for effortful listening, then the increased demands of CI-mediated speech perception may leave fewer resources available for generating predictions.
● **Audiovisual integration at the service of bottom-up integration rather than top-down prediction**. Although CI users are known to rely more heavily on visual speech information than NH listeners (Rouger et al., 2007; Stevenson et al., 2017; Strelnikov et al., 2009), the nature of this enhanced visual reliance may differ qualitatively from that of NH individuals. Specifically, CI users may use visual information primarily to support bottom-up speech recognition, disambiguating phonemes and segmenting the speech stream, rather than to facilitate top-down prediction. If visual speech cues are primarily allocated to recognition processes in CI users, they may not contribute to the generation of predictions in the same way they do in NH listeners. This interpretation suggests that visual speech information serves different functional roles depending on the integrity of the auditory system.
● **Strategic adaptation to unreliable predictions**. Given the variability and uncertainty inherent in CI-mediated perception, CI users may have learned that predictions are often unreliable guides to upcoming input. Such learning could lead to a strategic downregulation of predictive processing, a form of adaptation that prioritizes bottom-up evidence accumulation over top-down anticipation. This “wait-and-see” strategy has been proposed based on eye-tracking (Nagels et al., 2020) and pupillometry evidence (Winn, 2016) and would be adaptive in an environment where predictions frequently fail to match the input. The absence of pre-target oscillatory effects in CI users could reflect this strategic shift rather than an inability to generate predictions per se.

As a final note, when analyzed alone, CI users showed a power increase in the first half of the gap in the V-relative to the V+ condition. A visual inspection of the time-frequency representations (Figure 6) suggests that this increase may reflect an enhancement of alpha-band activity (8–11 Hz) during the transition from the sentence frame to the gap, an increase that seems particularly pronounced in the V-condition. Speculatively, this alpha increase may indicate a transient cortical disengagement when the mouth is occluded. Notably, CI users also exhibit stronger overall alpha synchronization than NH listeners, which may point to different strategies for integrating or compensating for visual information during speech processing. This enhanced alpha activity could reflect inhibitory processes that suppress irrelevant visual processing when articulatory cues are unavailable, or alternatively, could indicate reduced engagement of language networks when multimodal cues are incomplete. However, these observations remain purely speculative.

### 4.3 No post-target N400 modulation by group or mouth visibility

Despite the marked group differences in pre-target oscillatory activity, the N400 constraint effect was robust and comparable across CI users and NH controls. Both groups showed more positive ERPs to words in high-constraint relative to low-constraint contexts, indicative of facilitated lexical-semantic access and integration for predictable targets. The absence of a Group × Constraint interaction suggests that, once a word is encountered, CI users and NH listeners engage broadly similar mechanisms for evaluating the fit between the incoming word and the preceding context. This pattern converges with N400 studies in children with CIs, where semantic incongruity reliably elicits N400 effects that can approach or even exceed those of typically hearing peers, despite weaker behavioral semantic performance (Kallioinen et al., 2016; Pierotti et al., 2022; Vavatzanidis et al., 2018; Kallioinen et al., 2023). Together with our results, these findings indicate that the basic neural machinery supporting semantic access and integration, as indexed by the N400, can develop and remain functional in CI users despite long-term exposure to degraded auditory input.

Importantly, our CI sample consisted primarily of adolescents and young adults (mostly between 12 and 27 years) who had been implanted in early childhood (over half by age 5 and the majority before age 8) and had, on average, more than a decade of CI experience, alongside a smaller subset of older, late-implanted adults. This demographic profile places our participants in between the very young, early implanted children typically studied in developmental N400 work and the older, often postlingually deaf adults examined in many CI studies. Developmental work shows that earlier implantation and increasing auditory experience are associated with more robust and earlier N400 effects (Vavatzanidis et al., 2018; Kallioinen et al., 2023), whereas studies in older CI users and in adverse listening conditions often report reduced and delayed N400 predictability or congruency effects (e.g., Burkhardt et al., 2022; Hsin et al., 2025). Against this background, the preserved N400 constraint effect in our heterogeneous but predominantly early-implanted, long-term CI group suggests that once stable lexical–semantic networks have been established, CI users can exploit contextual constraints at the time of word processing to a degree comparable to NH listeners. At the same time, the inclusion of older, late-implanted participants likely increased variability in the CI group and would, if anything, bias against finding a robust N400 effect; the fact that we still observe one at the group level makes this conclusion conservative. Together with the pre-target group differences, this may hint at possible different mechanisms for CI users to represent and exploit context at the service of easier target processing.

The exploratory correlational analyses in the CI group (Supplementary Figures S7 and S8) suggest that individual differences in peripheral hearing may be are meaningfully related to electrophysiological outcomes. Within the CI group, poorer audiometric thresholds (higher PTA4) were associated with larger N400 constraint effects (LC–HC) in the target-locked ERP in both V+ and V-conditions (r ≈ –0.5). Although these correlations fell just short of conventional significance after FDR correction, they were consistently supported by multiple robustness checks, arguing against single-case or distribution-driven explanations. However, these correlations were not replicated when analyzing the 1.3-1.9 s window in the gap-onset ERP and should be treated as merely suggestive. No reliable associations emerged with years of CI use, either with target– or gap-locked ERPs. This pattern is compatible with prior mechanistic accounts suggesting that CI listeners may achieve comparable N400 profiles via different routes: in CI children, large N400 mismatch effects in long-SOA priming have been interpreted as stronger, top-down predictive strategies under sparse input (Kallioinen et al., 2016; Pierotti et al., 2022; Kallioinen et al., 2023), whereas in adults, both CI users and NH listeners under degraded or prolonged listening-predictability effects often attenuate and delay, consistent with less stable lexical expectations and slower integration (Burkhardt et al., 2022; Hsin et al., 2025). Our findings nuance this picture in a predominantly early-implanted, long-term CI sample: prediction error computation at word recognition appears relatively preserved (robust N400 constraint effect), but its magnitude scales with the degree of peripheral degradation, such that worse thresholds are linked to stronger contextual modulation, consistent with a compensatory shift toward semantic prediction when sensory evidence is limited. Coupled with the pre-target alpha–beta group differences we observed, the overall picture suggests that CI users operate under a chronically high “listening effort” regime that constrains how strongly and how early lexical predictions can be instantiated. The PTA4–N400 correlations further indicate that, within this regime, individuals with poorer auditory thresholds may compensate by generating stronger contextual predictions, which then manifest as larger N400 constraint effects at the target. Against this backdrop, the preserved N400 constraint effect in our sample likely reflects successful use of contextual information at the moment of word processing, even if predictive pre-activation during the gap (measured as beta power decrease) is absent in left frontal and central sensors, thus likely differently implemented and shaped by a variety of individual factors.

The three-way interaction observed in the exploratory analysis of the later time window (1.7–1.8 s post-gap onset) adds further nuance. In this window, CI users showed a larger constraint effect than NH controls specifically in the mouth-visible condition. CI users are known to exhibit enhanced lip-reading skills and to rely heavily on visual speech cues, and N400 work in children with CIs has been interpreted as evidence that they often lean on top-down semantic predictions when the signal is sparse or ambiguous. One plausible interpretation, particularly given our sample’s long CI experience and the PTA4–N400 relationships described above, is that when visual speech information is available during the sentence frame, CI users with poorer auditory thresholds build strong multimodal predictions that require extended post-lexical processing to reconcile with the acoustically degraded target. The prolonged constraint effect may thus index additional computational demands associated with integrating cross-modal predictions with impoverished auditory input, rather than a simple delay of the canonical N400.

We did not find clear effects of mouth visibility in the main analysis of the N400. This was less unexpected, as mouth visibility was manipulated only during sentence frame presentation, with the rationale of manipulating the informativity available to generate predictions (auditory only vs audiovisual). The lack of N400 effects is in line with work showing that facial cues mainly tune earlier phonological and lexical stages and late attention, while semantic integration itself is relatively robust to mouth visibility manipulations. In Hernández-Gutiérrez et al. (2018), dynamic vs static faces and selective occlusion of the mouth or eyes were manipulated while participants listened to constraining sentences; across three experiments, the N400 to unexpected words was remarkably insensitive to all facial-feature manipulations, even though dynamic faces reliably elicited a late posterior positivity for expected words and were interpreted as increasing motivated attention in easy conditions. This fits with broader audiovisual evidence: visual speech typically speeds and sharpens early auditory processing (reduced and earlier N1/P2) and improves intelligibility, particularly in noise, without necessarily changing the size of the semantic N400 effect when the auditory signal is already clear (van Wassenhove et al., 2005). When N400 modulations do appear, they tend to arise under conditions where visual articulatory information is highly diagnostic about the upcoming word (high saliency) or when the acoustic input is degraded, and they are often interpreted as late phonological/lexical influences rather than changes in core semantic integration (Brunellière et al., 2013). In our design, mouth visibility differed only during the sentence frame, but the critical word was always accompanied by visible articulation in both conditions, and the acoustic signal at the target was clear and informative. Under these circumstances, visual speech during the frame might help set up lexical and phonological predictions or reduce processing effort, but at target onset both conditions converge on very similar audiovisual input; lexical fit is computed on essentially the same evidence, so the N400 constraint effect remains unchanged. Finally, the general delay in the emergence of N400 relative to typical auditory paradigms deserves mention (Kutas & Van Petten, 1994). This shift is explained by variability in the interval between the onset of the video recording (which starts immediately after the silent gap at 0.8 s) and the actual onset of the spoken word in each recording. In other words, there is non-negligible jitter between the beginning of the video and the moment in which the auditory stimulus and the corresponding articulatory movements begin. This variability is further amplified by the fact that the onsets of voicing and mouth movements do not follow a fixed sequence but depend on the phonetic properties of the upcoming word. For example, voiceless stop consonants begin with visible articulatory movements, with voicing emerging only at the onset of the following vowel; in contrast, voiced stops begin with vocal-fold vibration before the lips reach the occlusion position. As a result, the temporal alignment between video onset, mouth movement onset, and voice onset varies considerably across items. On average, this delay corresponds to 219 ms (SD = 81.1) for voice onset and 165 ms (SD = 141) for mouth movement onset (Supplementary Figure S4). When these delays are added to the target onset at t = 0, the later emergence of the N400 becomes expected (see also the voice-locked ERP in Supplementary Figure S4, which clearly displays that when voice onset is taken as t = 0 the N400 emerges as early as around 200 ms, as expected for auditory words).

### 4.4 Prediction generation is linked to language production skills

The robust negative correlation between pre-target beta power decreases and language production skills provides important evidence for theories linking prediction and production mechanisms. Specifically, individuals with higher production proficiency, as indexed by a composite of verbal fluency and sentence generation performance extracted via PCA, showed larger constraint-related power decreases over left frontal and central sensors. This finding supports the hypothesis that prediction during comprehension involves covert simulation of upcoming speech using the language production system (Federmeier, 2007; Gastaldon et al., 2024a; Pickering & Garrod, 2013; Pickering & Gambi, 2018; Pickering & Strijkers, 2025). The prediction-by-production account proposes that comprehenders generate predictions by running their production system “in reverse,” using contextual information to internally simulate likely upcoming words or phrases. The robust left-lateralized frontal distribution of the correlation effect we found is consistent with the involvement of inferior frontal regions in both speech production and predictive language processing (Hagoort, 2013; Wang et al., 2018). This finding has particular relevance for understanding and potentially improving language processing in CI users. If prediction relies on production mechanisms, and if CI users show weaker predictive oscillatory signatures alongside lower production scores, then interventions targeting expressive language skills might have downstream benefits for receptive processing. Speech-language therapy for CI users has traditionally emphasized perception and comprehension; our results suggest that strengthening production abilities could enhance the predictive mechanisms that support efficient comprehension. This hypothesis warrants direct testing in intervention studies, as the correlational nature of our data precludes strong causal claims, and alternative interpretations should be considered.

### 4.5 Clinical and theoretical implications

The present findings have several implications for clinical practice and theoretical models of language processing under adverse conditions. From a clinical perspective, the absence of pre-target predictive signatures in CI users, despite preserved post-target integration effects, suggests that rehabilitation approaches might beneficially target the generation phase of prediction. Encouraging CI users to actively engage in anticipatory mechanisms, for instance through explicit instruction in using contextual cues or through training paradigms that reinforce prediction, could potentially enhance comprehension efficiency and reduce listening effort. The strong correlation between production skills and predictive processing supports the inclusion of expressive language training in comprehensive CI rehabilitation programs. While current approaches emphasize auditory training and speech perception, our findings suggest that developing robust production abilities may have cascading benefits for comprehension by strengthening the prediction mechanisms that support efficient language processing.

From a theoretical standpoint, our results contribute to ongoing debates about the nature and variability of predictive processing. The data suggests that prediction is not an all-or-nothing phenomenon but rather a flexible mechanism that varies across individuals and is modulated by factors including sensory input quality, available cognitive resources, and the presence of complementary information sources (Gastaldon et al., 2024a). The emergence of prediction-related oscillatory effects specifically in the mouth visible condition for NH controls suggests that the integration of meaningful multimodal speech cues plays a facilitatory role in prediction generation. This finding extends previous work showing that audiovisual speech benefits exceed simple signal enhancement (Peelle & Sommers, 2015) by demonstrating that visual speech information specifically enhances anticipatory neural processes. Models of audiovisual speech processing may need to incorporate this predictive dimension to fully account for the multimodal advantage in speech comprehension.

### 4.6 Limitations and future directions

Several limitations of the present study should be acknowledged. First, the sample size (N = 18 per group), while typical for EEG studies with clinical populations, limits statistical power for detecting smaller effects and interactions. The absence of certain effects (e.g., the constraint × mouth visibility interaction in CI users) should be interpreted cautiously, as null findings may reflect insufficient power rather than true absence of effects. Replication with larger samples is warranted. Such a limitation could be overcome by multi-lab studies (Heinrich & Knight, 2020; Lange, 2020; McShane et al., 2019). Second, the CI sample was heterogeneous in terms of age of implantation, duration of CI use, etiology of deafness, and implant configuration (unilateral vs. bilateral). While this heterogeneity reflects the clinical reality of CI populations, it introduces variability that may obscure systematic effects. Future studies with more homogeneous samples, for instance, focusing specifically on early-implanted adults or sequential bilateral CI users, could provide clearer insights into how specific factors modulate predictive processing. Third, the electrode placement limitations imposed by the cochlear implant precluded full coverage of temporo-parietal regions in CI users, creating a baseline imbalance in data collection across groups. While we mitigated this issue by excluding temporal ROIs from between-group comparisons, this limitation reduces our ability to characterize the full spatial distribution of effects in CI users and to directly compare temporo-parietal activity across groups, preventing us from assessing potential effects in sensors with unavailable data. We cannot exclude, in fact, that some contextual representation was formed in temporo-parietal regions, thus explaining more easily the presence of a N400 in lack of pre-target effects in the available sensors we analyzed. Anecdotally, by observing the left temporo-parietal ROI in the CI users with right-ear implant (N = 6), this might be the case. However, this remains highly speculative. Finally, we did not assess cognitive factors such as working memory capacity or processing speed that might modulate predictive processing independently of hearing status. Including such measures in future studies would help disentangle sensory from cognitive contributions to the observed group differences.

Future research should explore several directions suggested by the present findings. Longitudinal studies following CI users from implantation could reveal how predictive processing develops or changes with auditory experience. Despite the challenges imposed on neuroimaging techniques by the implant, future technical improvements could help identify the neural generators of prediction-related effects and clarify whether CI users recruit partly different networks for language processing. Intervention studies testing whether production training enhances prediction would provide causal evidence for the production-prediction link. Finally, studies comparing different visual manipulations (e.g., lips only, full face, static vs. dynamic) could clarify which aspects of visual speech information most effectively support prediction.

## 5. Conclusions

The present study provides the first EEG evidence on predictive processing during audiovisual speech comprehension in cochlear implant users. Our findings reveal a complex picture in which pre-target neural signatures of prediction generation are altered in CI users while post-target integration processes remain intact and comparable to those of NH listeners. The emergence of prediction-related time-frequency effects specifically when mouth cues were available, and only in NH controls, highlights the facilitatory role of visual speech information in supporting anticipatory language processing. The correlation between production abilities and prediction strength provides support for production-based models of prediction and suggests potential avenues for enhancing language processing in CI users through expressive language training. Together, these findings advance our understanding of how sensory limitations shape the neural mechanisms of language comprehension and open new directions for both basic research and clinical intervention.

## CRediT author statement

**Simone Gastaldon**: Conceptualization, Methodology, Software, Validation, Formal analysis, Investigation, Data Curation, Writing – Original Draft, Visualization, Funding acquisition. **Flavia Gheller**: Conceptualization, Methodology, Investigation, Writing – Original Draft. **Noemi Bonfiglio**: Investigation, Writing – Review and editing. **Davide Brotto**: Methodology, Resources, Writing – Review and editing. **Davide Bottari**: Methodology, Writing – Review and editing.

**Patrizia Trevisi**: Methodology, Resources, Writing – Review and editing. **Alessandro Martini**: Methodology, Resources, Writing – Review and editing. **Francesco Vespignani**: Conceptualization, Methodology, Software, Writing – Review and editing. **Francesca Peressotti**: Conceptualization, Methodology, Supervision, Writing – Review and editing, Project administration, Funding acquisition.

## Conflict of interest

None

## Data availability statement

Raw and preprocessed data with scripts necessary to reproduce the main findings of the study can be found in the Open Science Framework repository at the following URL: https://osf.io/g2kbj/overview?view_only=3606247f01524feb9d2e60d0c364797e

## Declaration of generative AI and AI-assisted technologies in the manuscript preparation process

During the preparation of this work the authors used ChatGPT and Claude AI in order to make code more efficient and to improve manuscript grammar and readability. After using these tools, the authors reviewed and edited the content as needed and take full responsibility for the content of the published article. No content was generated with the sole use of generative AI. ChatGPT was used to generate the human face used in Figure 2, exclusively for privacy reasons and descriptive purposes.

## Funding

This research was supported by the Italian Ministry of University and Research with the PRIN grant for the project “The role of cochlear implantation and bimodal bilingualism in early deafness: a window into the neurofunctional mechanisms of human language” (project code 20177894ZH), awarded to F.P. S.G. received support from a postdoctoral fellowship funded through the aforementioned PRIN grant (2021-2023), and from a fellowship funded by Fondazione CARIPARO (PHD@UNIPD) for the project “Predictive Brain in Audiovisual Speech Comprehension” (CUP_C93C23003190005), awarded to S.G (2023-2025).

## Acknowledgements

We thank all participants and their caregivers for their time and willingness to take part in the study. We are also grateful to Bianca Bonato for serving as the on-camera speaker for the recorded sentence stimuli, and to Eleonora Sofia Fazio and Marta Battistutta for their assistance with data collection and scoring.

## Supplementary Figures

**Figure S1:**
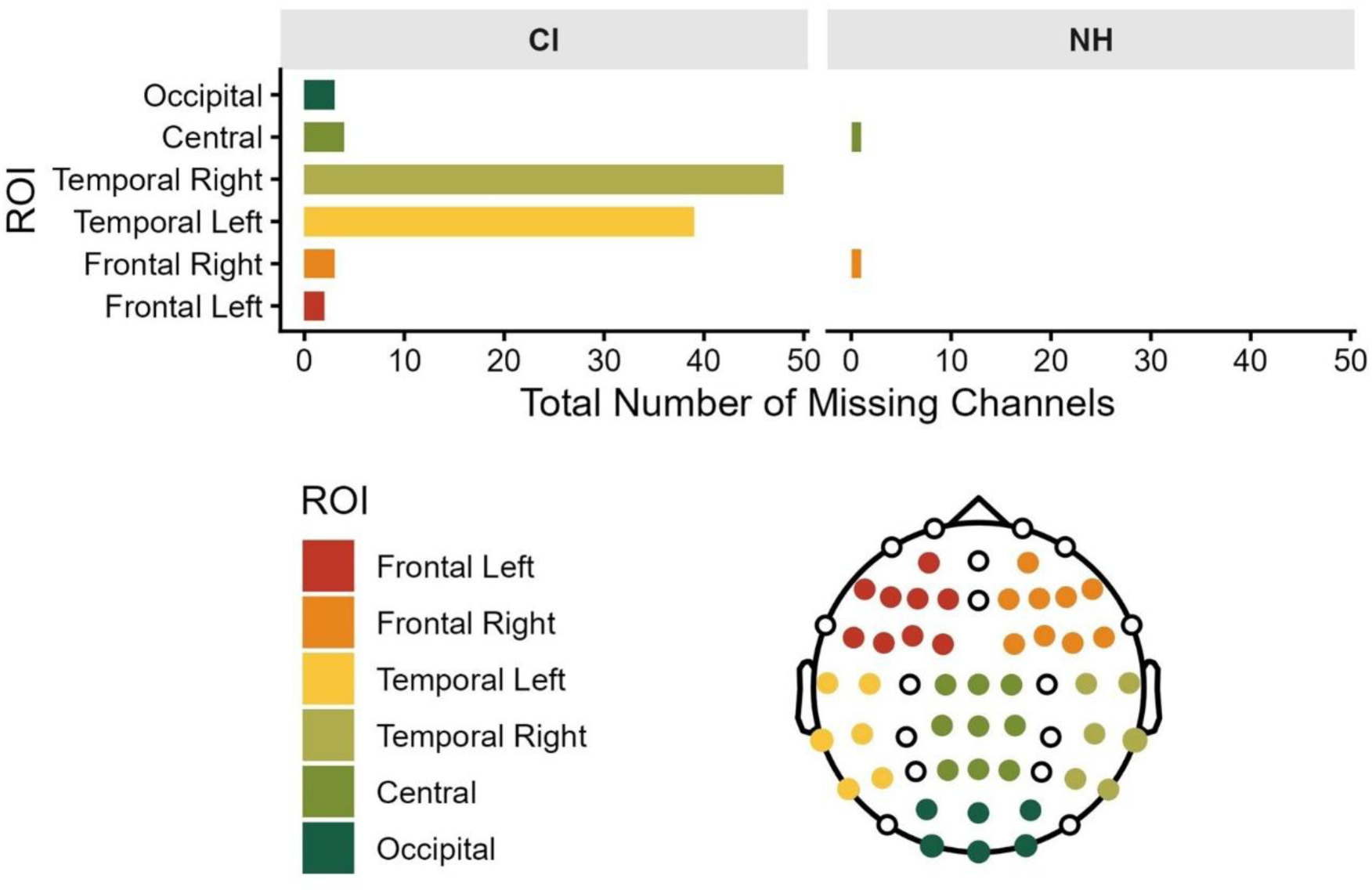
Regions of interest and missing channels. For each ROI as defined in the Methods section, the number of missing channels belonging to each ROI in each group is plotted. The CI group misses many temporal channels and therefore analyses on such ROIs are not performed when including CI users.

**Figure S2:**
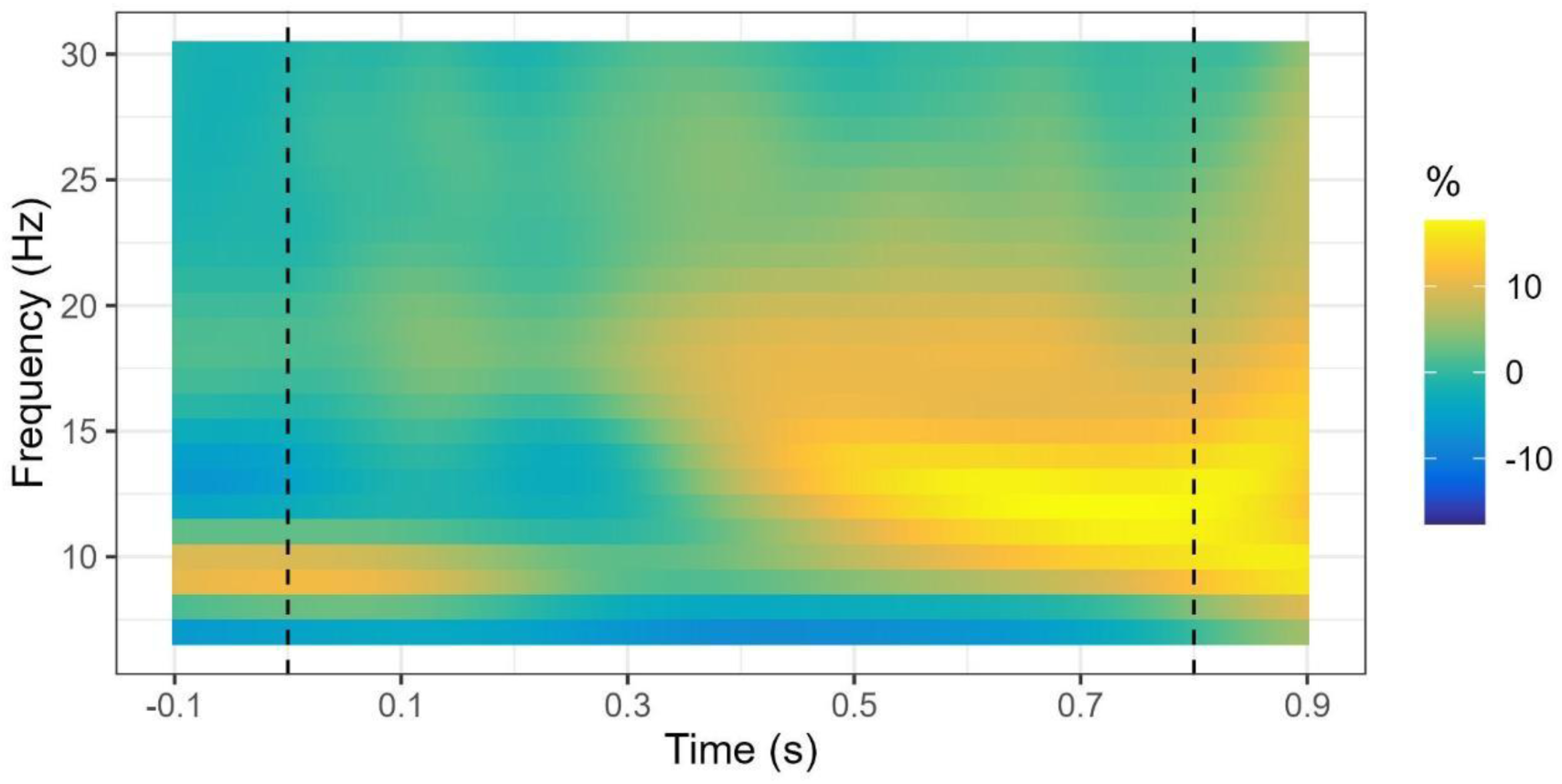
Time-frequency grand-average at the gap. All participants and conditions are averaged together to determine the overall modulations across frequency and time. This allowed us to identify the two time-windows of the gap (0-0.4 s and 0.4-0.8 s) and the four frequency ranges of interest (8-11 Hz, 12-15 Hz, 16-20 Hz, 21-30 Hz).

**Figure S3:**
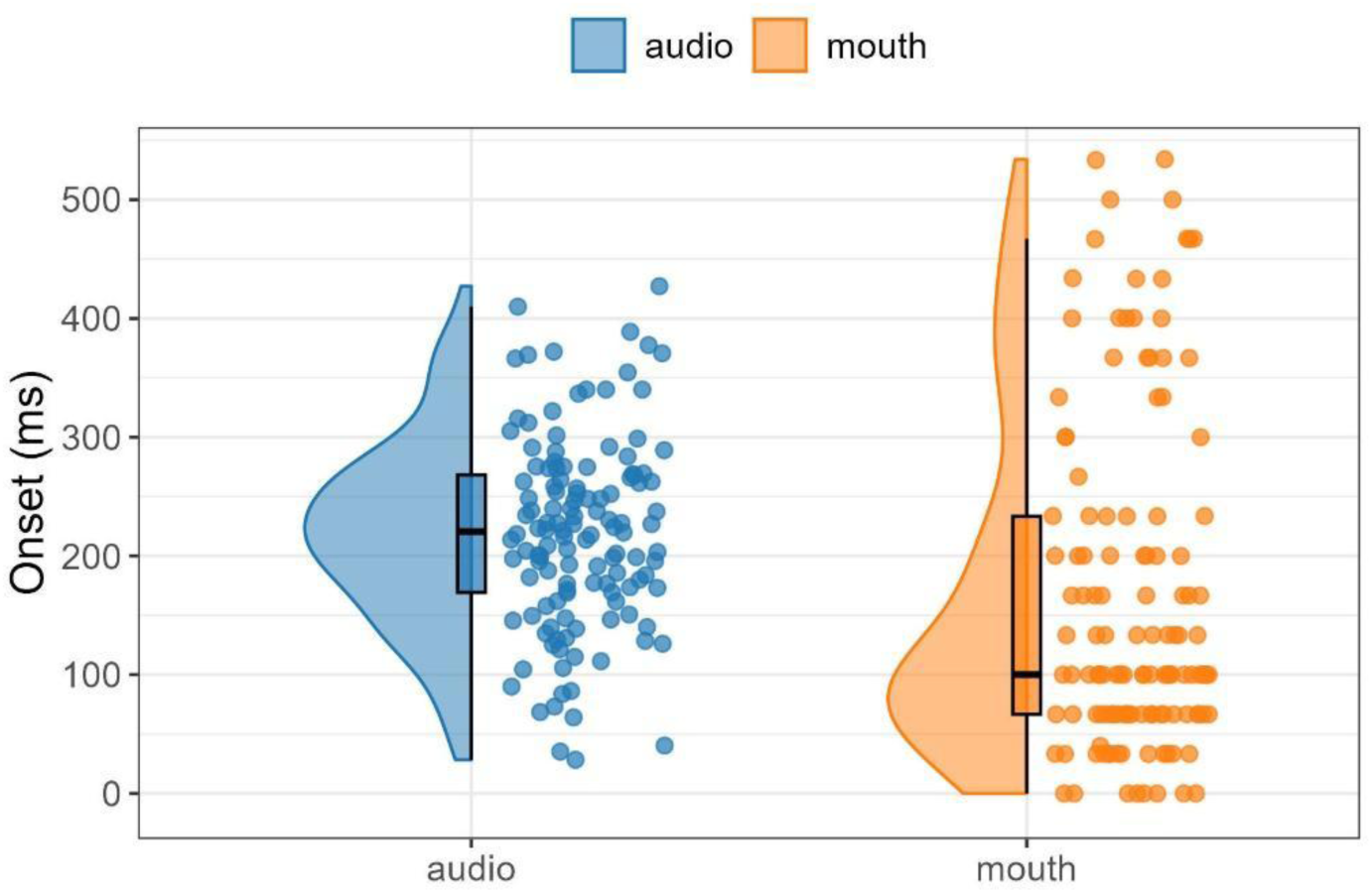
Delays between file onset and speech and mouth movement onset. The two distributions show that between the onset of the video (t_0_ = target onset) and the actual onset of the audio and the mouth movements of each item is variable. There is an inherent variable jitter between the onset of the file and the actual onset of the stimuli, variable across sound onset. This explains the delay in the emergence of the N400.

**Figure S4:**
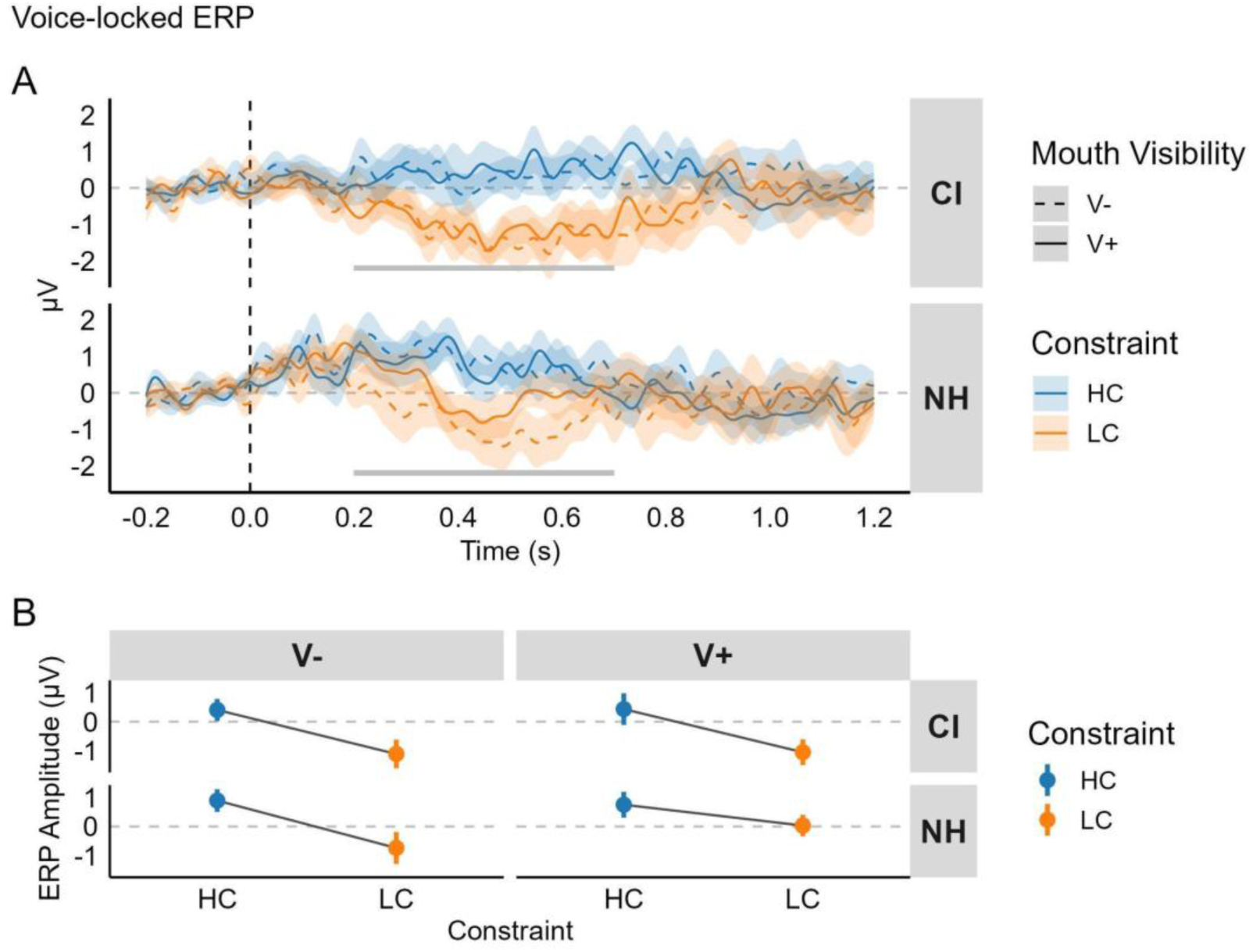
ERPs computed at the onset of the actual voice onset. The time-series show that the artifactual interaction in NH listeners is still present (LC_V– being more negative). It also shows, however, that the N400 emerges more clearly and temporally in line with the expectations when t_0_ is aligned with speech onset.

**Figure S5:**
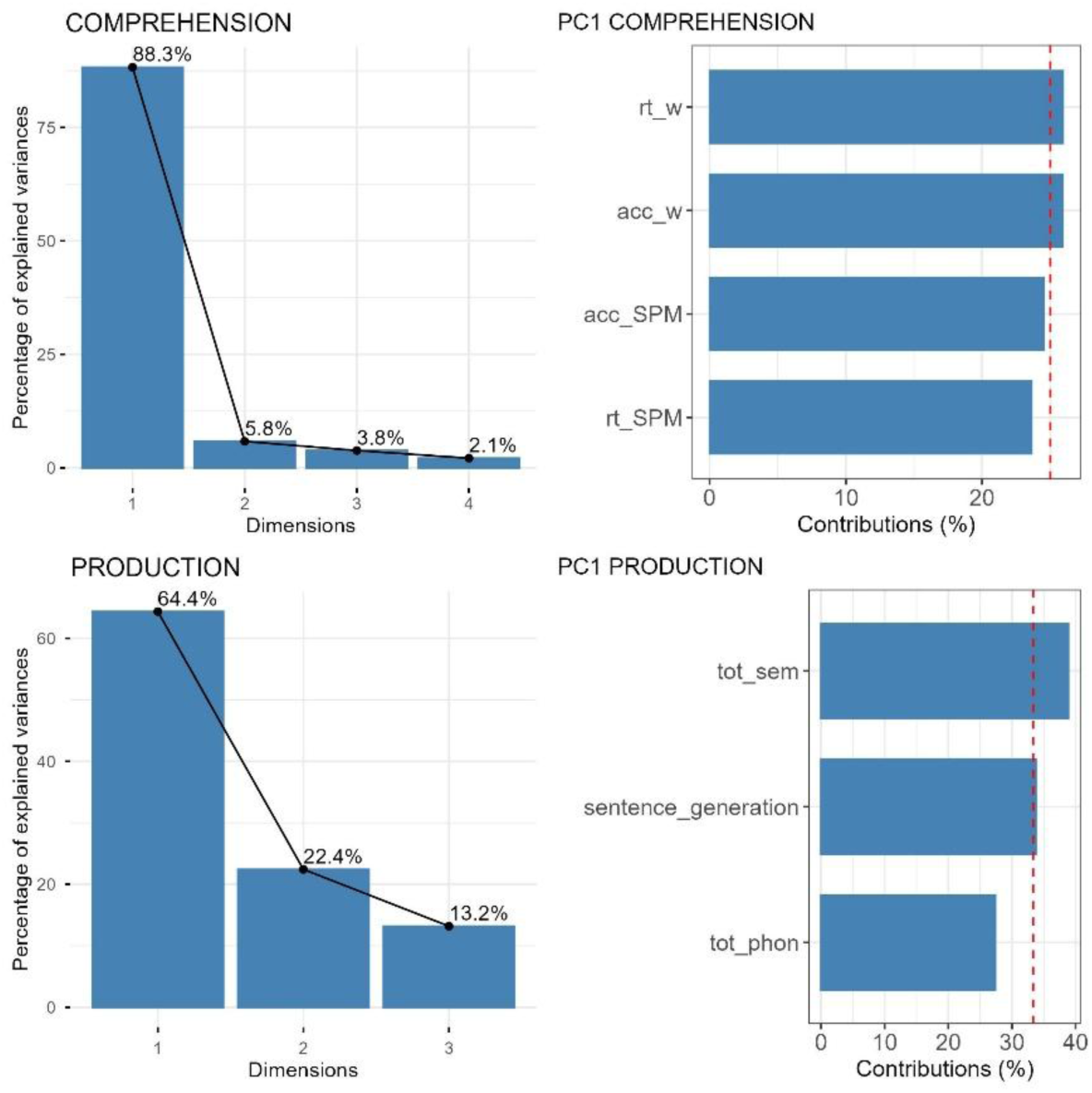
PCA loadings for the comprehension and production language scores. For each task sets (comprehension and production), the figure shows the total variance explained by the principal components is shown (left), and the % contribution of each measure to the first component (PC1; right).

**Figure S6:**
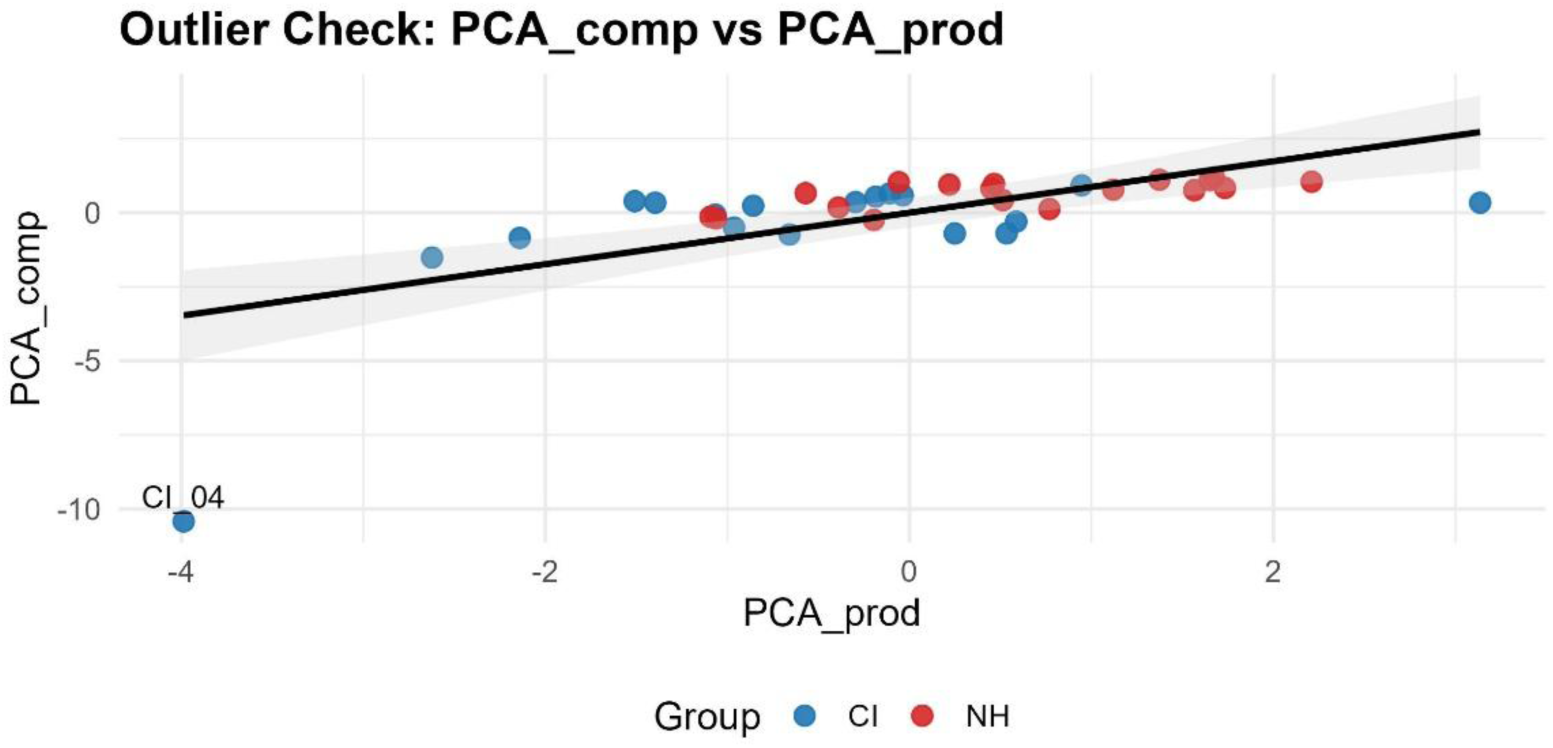
Outlier in the PC1 comprehension. One participant of the CI group (CI_04) resulted to be an outlier in the PC1 of the comprehension and production scores. This participant has been removed from the correlations with PC comp and prod.

**Figure S7.**
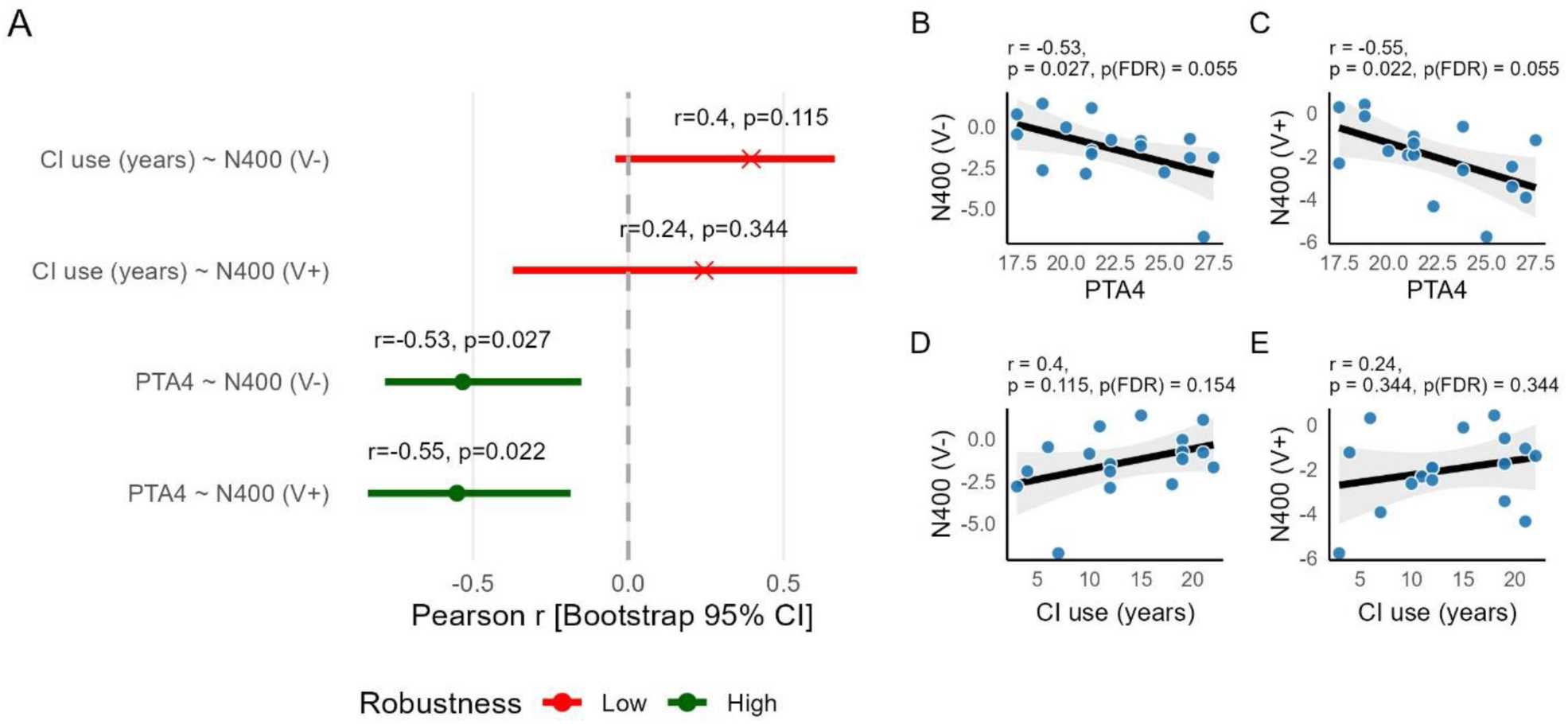
Robustness check on N400-audiometric and CI use correlations (target-locked; 0.6-1 s). A) Result of the robustness check procedure. Color represents the composite score describing correlation robustness. B-E) Scatterplots for each correlation between Years of CI use and PTA4 and the N400 effect in the two mouth visibility conditions.

**Figure S8:**
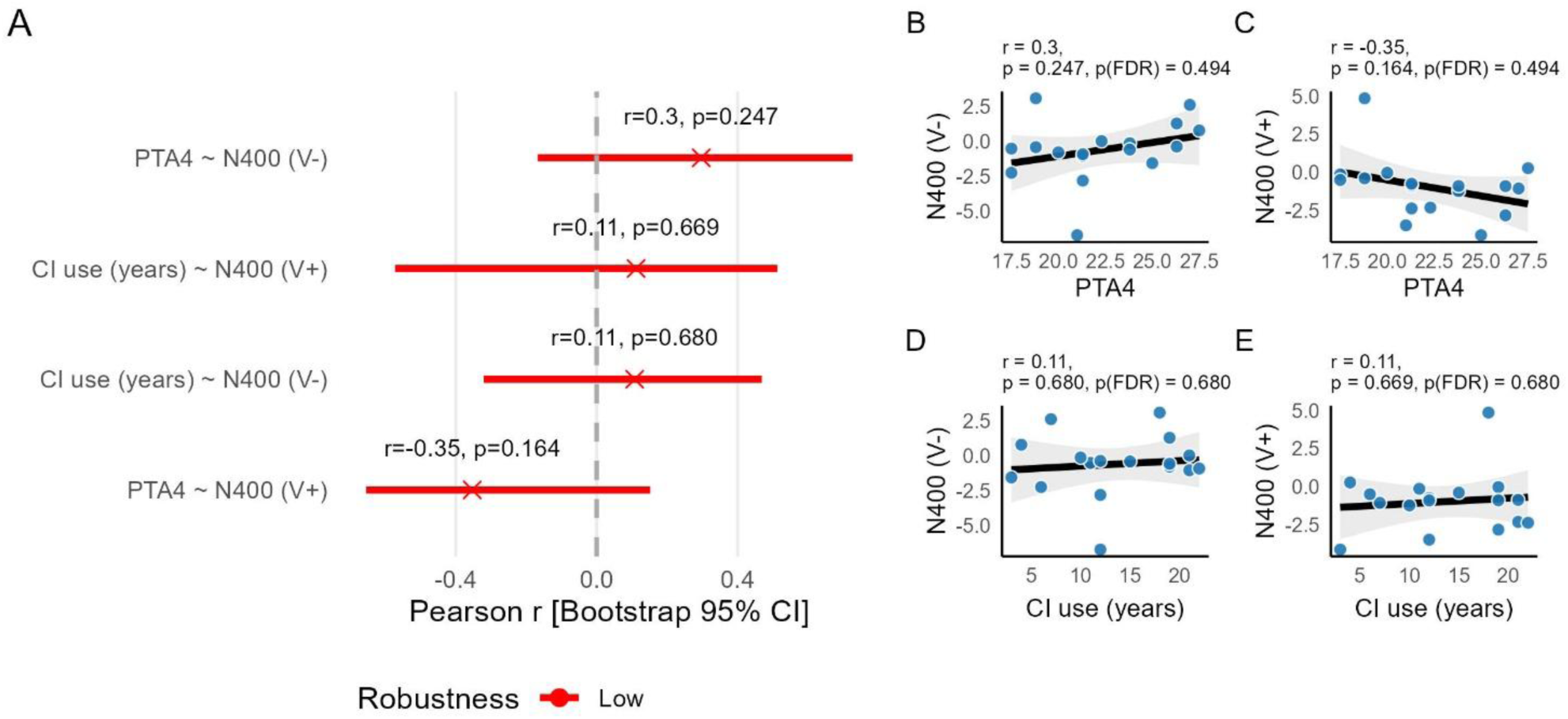
Robustness check on N400-audiometric and CI use correlations (gap-locked; 1.4-1.9 s). A) Result of the robustness check procedure. Color represents the composite score describing correlation robustness. B-E) Scatterplots for each correlation between Years of CI use and PTA4 and the N400 effect in the two mouth visibility conditions.

## Footnotes

1 https://ripi.iss.it/ripi/it/il-progetto/ridiu-registro-italiano-dispositivi-impiantabili-uditivi/

## Notes

https://osf.io/g2kbj/overview?view_only=3606247f01524feb9d2e60d0c364797e

### Competing Interest Statement

The authors have declared no competing interest.

